# Predicting intelligence from fMRI data of the human brain in a few minutes of scan time

**DOI:** 10.1101/2021.03.18.435935

**Authors:** Gabriele Lohmann, Eric Lacosse, Thomas Ethofer, Vinod J. Kumar, Klaus Scheffler, Jürgen Jost

## Abstract

In recent years, the prediction of individual behaviour from the fMRI-based functional connectome has become a major focus of research. The motivation behind this research is to find generalizable neuromarkers of cognitive functions. However, insufficient prediction accuracies and long scan time requirements are still unsolved issues. Here we propose a new machine learning algorithm for predicting intelligence scores of healthy human subjects from resting state (rsfMRI) or task-based fMRI (tfMRI). In a cohort of 390 unrelated test subjects of the Human Connectome Project, we found correlations between the observed and the predicted general intelligence of more than 50 percent in tfMRI, and of around 59 percent when results from two tasks are combined. Surprisingly, we found that the tfMRI data were significantly more predictive of intelligence than rsfMRI even though they were acquired at much shorter scan times (approximately 10 minutes versus 1 hour). Existing methods that we investigated in a benchmark comparison underperformed on tfMRI data and produced prediction accuracies well below our results. Our proposed algorithm differs from existing methods in that it achieves dimensionality reduction via ensemble learning and partial least squares regression rather than via brain parcellations or ICA decompositions. In addition, it introduces Ricci-Forman curvature as a novel type of edge weight.

## Introduction

Many neurological and psychiatric conditions evade detection by standard anatomical MRI. There is a growing hope that functional MRI (fMRI) may help to fill this gap. In this context, the suitability of fMRI for predicting individual behaviour has been investigated in a number of recent studies [1–10]. Predictive modelling generally proceeds in three stages [11]. First, a dimensionality reduction via brain parcellations or ICA decomposition is performed. Second, interactions between the brain parcels or components are estimated. Finally, a classifier or regressor is trained to predict behavioural traits or other quantities of interest. The various methods differ with respect to the choice of the strategies used in those three stages, but some form of network modelling is common to all [12, 13]. The methodological challenges and best practices are discussed in [14–18].

Here we introduce a new machine learning algorithm for predicting intelligence from fMRI, and validate it on data of the Human Connectome Project (HCP) [19, 20]. Specifically, we investigate the predictability of general intelligence as defined by Dubois et al. [21], and of fluid, crystalline, and total intelligence as defined in the behavioural testing protocol of the Human Connectome Project [22]. Human intelligence and its neural representations have been a focus of research for many years, see e.g. [23–26]. However, our focus here is on the machine learning methodology of intelligence prediction rather than on the topic of intelligence in itself. Thus, our proposed algorithm is not intended to be limited to the prediction of intelligence. Rather, we view this as a proof of concept for a wider range of potential applications [27, 28].

We call our algorithm VEGA (**V**ox**E**l-**G**raph machine learning **A**lgorithm). VEGA differs in several respects from the predictive modelling strategy that is generally adopted in other studies. First, it does not perform brain parcellations or ICA decompositions for dimensionality reduction. Instead, it employs ensemble learning and partial least squares regression to deal with the high dimensionality of the data. We also use the novel concept of Ricci-Forman curvature to define a brain mask to further constrain dimensionality. Second, VEGA works directly in voxel space, it does not require a surface extraction as some other methods do. This may be advantageous in situations where data quality is poor so that image segmentations and surface extractions may be problematic. This is particularly relevant in clinical environments where data quality can be an issue. And finally, VEGA is designed to work on both resting state (rsfMRI) as well as on task-based fMRI (tfMRI). We will show that its performance on tfMRI is particularly encouraging even though the scan times were very short.

### Experimental data and preprocessing

We downloaded fMRI data of 390 unrelated subjects (202 female, 188 male, aged 22-36, median age 28) acquired at 3 Tesla by the Human Connectome Project (HCP), WU-Minn Consortium [19,20,29]. Only subjects for whom all necessary data sets, i.e. rsfMRI, tfMRI and intelligence test scores were complete and fully available were included. We excluded data sets for which data quality problems due to instability of the head coil were reported [30]. All subjects gave written, informed consent according to the protocol by the Human Connectome Project consortium.

The rsfMRI data were acquired in two sessions on two separate days with two different phase encoding directions (left-right and right-left) with spatial resolution (2*mm*)^3^, multiband factor 8.

Each scan hat 1200 volumes acquired at TR=0.72 seconds so that the total scan time across all four sessions was approximately 58 minutes. The rsfMRI data were minimally preprocessed and cleaned using FSL-FIX [31–33].

In addition, we downloaded minimally preprocessed tfMRI data from two tasks, namely the language task and the working memory task [29]. The working memory task followed an N-back paradigm where participants were presented with pictures of places, tools, faces and body parts. The language paradigm consisted of a story comprehension task interleaved with a math task, see [34]. We chose those two tasks because they appear to be more closely linked to intelligence than any of the other tasks included in HCP. The tfMRI data were acquired in two sessions each (left-right and right-left phase encodings). The language task had 316 volumes per session (total scan time ≈ 7.5 min). The working memory task had 405 volumes per session (total scan time ≈ 10 min).

The tfMRI data were additionally subjected to a temporal highpass-filter (cutoff frequency 1/100 Hz) to remove baseline drifts. For both rsfMRI and tfMRI data, we reduced the spatial resolution to (3*mm*)^3^ via trilinear interpolation to limit the computational load. Furthermore, to counteract intersubject anatomical variability, we applied a spatial Gaussian filter using fwhm=6mm.

### Measures of Intelligence

Here we focused on four measures of intelligence that resulted from cognitive tests performed by the Human Connectome Project (HCP). For ease of notation, we use the same acronyms as in HCP to denote the various cognitive tests, see [22, 35]. The first two measures are fluid cognition (CogFluidUnadj) and crystallized cognition (CogCrystalUnadj). They are defined via averaging normalized scores of several tests as shown in table 1. The third measure (CogTotalUnadj) is a combination of the first two. The fourth measure is a general intelligence score (G-factor). Here we closely followed the work by Dubois et al. [21] who used a weighted average of normalized scores of a wider range of cognitive tests. We used the same test scores and the same weights for the average.

**Table 1:**
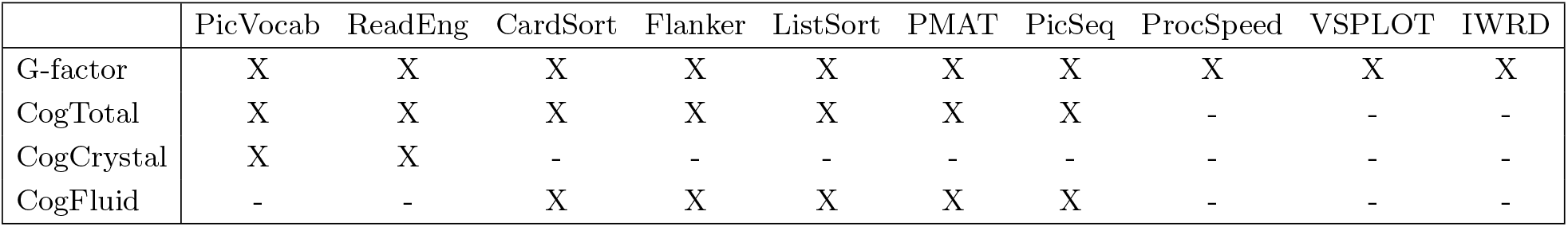
Components of the four measures of intelligence. The four measures result from averaging across the normalized test scores of the cognitive tests listed in this table. The acronyms are as in HCP [35]. For the G-factor, we used a weighted average of the ten test scores where the weights are as in [21], i.e. 0.624, 0.642, 0.364, 0.259, 0.451, 0.626, 0.354, 0.232, 0.578, 0.294 (in the same order as in the table).

#### Potential confounds

As in [21] we used multiple linear regression to regress out several potential confounds from the intelligence scores, namely handedness, gender, age (*Age_in_Yrs*), brain size (*FS_BrainSeg_Vol*) and the multiband reconstruction algorithm (*xf MRI_*3*T_ReconV rs*). To avoid leakage from training to test, the multiple linear regression was fitted on the training data, and the resulting weights were then used to remove the confounds in both training and test.

### A new algorithm for predicting intelligence from fMRI

We propose a new algorithm called “VEGA” for predicting intelligence from fMRI data of the human brain. It consists of four steps that are described in the following. For an overview, see Figure 1.

**Figure 1:**
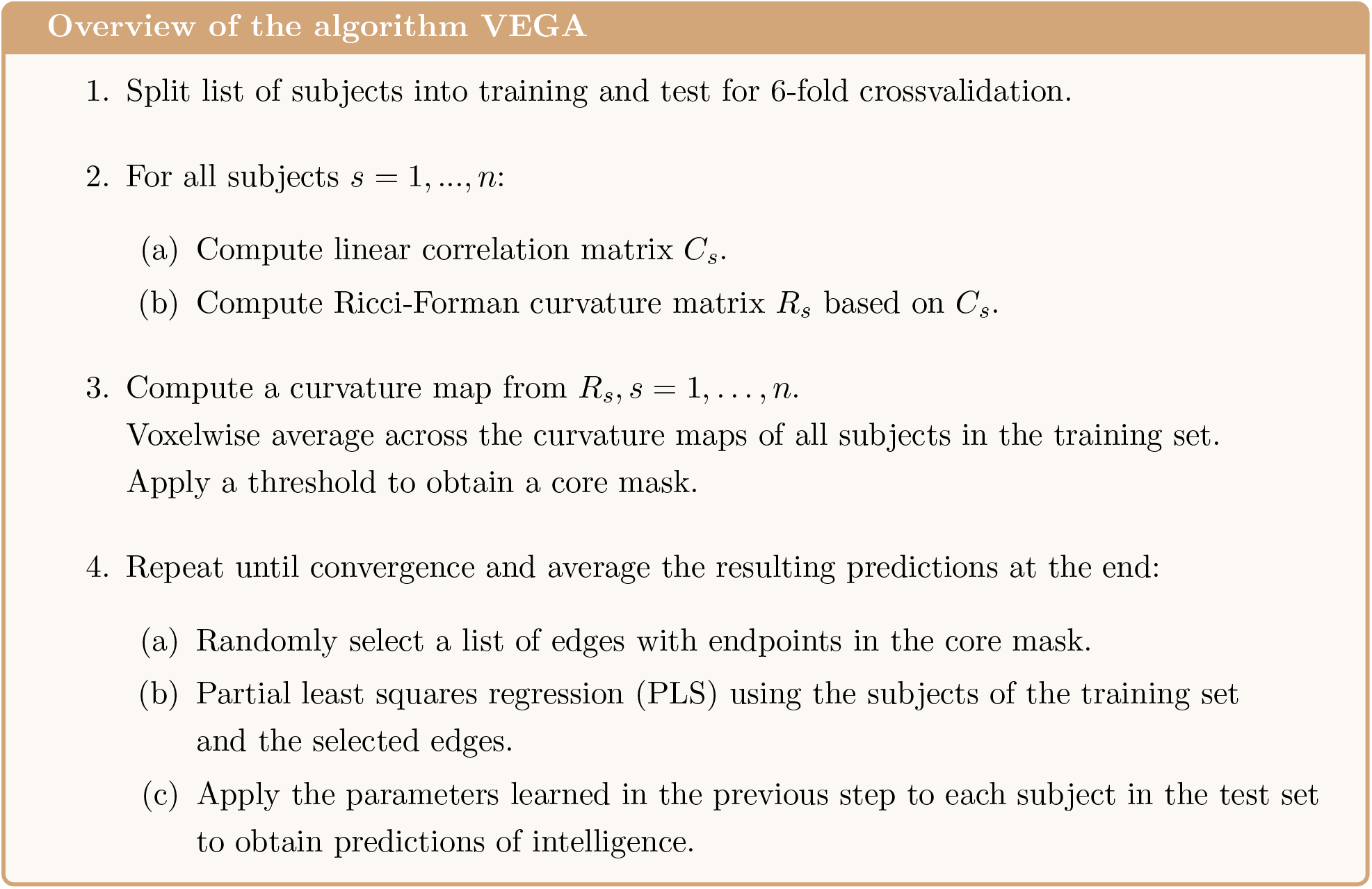
Overview of the algorithm. The proposed algorithm VEGA consists of the four steps as depicted in this overview. A main difference to existing algorithm is that it handles high-dimensionality via ensemble learning, partial least squares and Ricci-Forman curvature maps rather than via brain parcellations or ICA decompositions. A detailed description of each step is given in the main text.

#### Step 1 (Crossvalidation)

The set of *n* = 390 subjects is randomly split into six folds for crossvalidation. In each fold, 325 subjects are used for training a regression model that is subsequently tested on the remaining 65 subjects. This procedure is repeated for each of the six folds so that every subject is tested exactly once. In our experiments, we used twenty different and randomly selected train/test splits.

#### Step 2 (Connectivity matrices)

For each subject *s* = 1, …, *n* symmetric connectivity matrices *C*_*s*_, *R*_*s*_ are computed using two different measures of connectivity. Both matrices are of dimension *k* × *k* where *k* denotes the number of voxels in a brain mask. In the experiments reported below, the mask covers the entire brain with *k* = 55, 856 voxels, see Supplementary Figure 1.

#### Linear correlation

The first connectivity matrix *C*_*s*_ is based on the linear Pearson correlation coefficient. Its elements *c*_*i,j*_, *i, j* = 1, …*k* record the linear correlation between the fMRI time series in voxels *i* and *j* where *k* is the number of voxels in the brain mask. The matrix *C*_*s*_ is initially dense, i.e. it is computed for every pair of voxels in the brain mask so that the number of edges is *k*(*k* − 1)*/*2 ≈ 1.6^9^. From this large set of edges, we randomly select 1 million edges to which all subsequent analysis steps involving *C*_*s*_, *s* = 1, …, *n* are restricted. The purpose of this step is to reduce the memory load and computational burden. Note that we preselect edges, not voxels. Since the number of preselected edges is very large, all voxels in the brain mask serve at least once as an endpoint to one of those edges. This strategy provides a dense coverage of the brain that allows us to visualize brain areas that are predictive of intelligence (Supplementary Figure 3).

### Ricci-Forman curvature

Using the dense correlation matrix *C*_*s*_ computed in the previous step, a second connectivity matrix *R*_*s*_ is computed for every subject *s* = 1, …, *n*. It is based on the novel concept of Ricci-Forman curvature [36–38]. The motivation to apply this concept here is that it allows to attribute weights to edges that reflect their importance for the cohesiveness of a graph, and thus help to identify edges that are more reliable predictors. Ricci-Forman curvature has been previously applied to fMRI data [39], and more recently to diffusion weighted imaging (DWI) [40]. It defines a curvature for an edge *e* in a set of edges *E* as follows. Let *e* = (*v*_1_, *v*_2_) ∈ *E* and 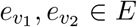 be any edges adjacent to *e* at vertices *v*_1_ and *v*_2_ with edge weights *ω*(*e*) and node weights *ω*(*v*). Then the Ricci-Forman curvature Ric_*F*_ of edge *e* is defined as

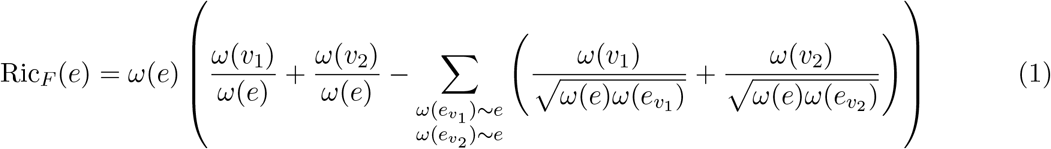

Figure 2 illustrates the geometric intuition behind this concept. Edges connecting vertices of large degree have strongly negative curvature values and may be interpreted as being most important for the cohesion of the network. For more information about the theoretical background of Ricci Forman curvature see [37, 38, 41].

**Figure 2:**
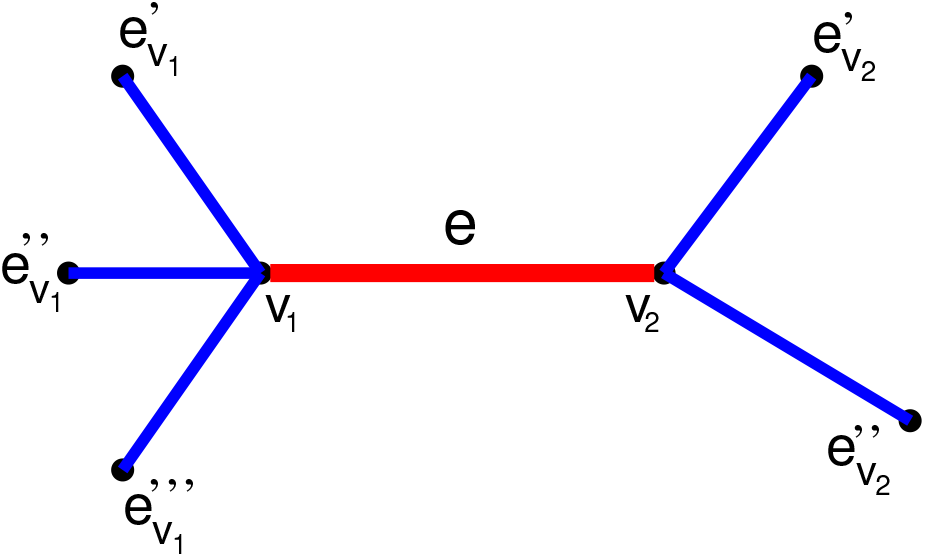
Illustration of Ricci-Forman curvature. Edge e with adjacent vertices v_1_ and v_2_ and parallel edges 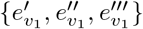 (adjacent to v_1_) and 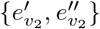 (adjacent to v_2_). The Ricci-Forman curvature of edge e is strongly negative if the edges parallel to its endpoints have large weights.

In the present context, vertices correspond to voxels and have a constant weight of 1. The edges correspond to correlations between fMRI time series of those voxels. From equation 1 we see that the edge weights *ω*(*e*) must be positive so that the Pearson linear correlation coefficient cannot be directly used as an edge weight. We therefore define an edge weight function *ω* applied to the correlation coefficient *x* as *ω*(*x*) = max(*x, ε*) where *ε* is a small positive constant. In our experiments, this constant was set to *ε* = 1*/k* where *k* is the number of voxels in the brain mask. With these definitions, we now compute a dense Ricci curvature matrix *R*_*s*_ for every subject *s* = 1, …, *n* using the corresponding correlation matrix *C*_*s*_ as input where each correlation value is first transformed via the weight function defined above. Note that the matrix *R*_*s*_ is symmetric and has *k*(*k* − 1)*/*2 distinct edges.

### Step 3 (Core mask)

Edges with strongly negative Ricci-Forman curvatures may be viewed as backbones of the correlational structure in the brain. We hypothesize that such edges may be particularly robust features for predicting intelligence. Here we use this idea to derive a new brain mask to which all subsequent analysis steps are constrained. This mask is obtained as follows.

First, we compute a “curvature map” separately for every subject. This is done by projecting all edges of *R*_*s*_ to a map in which a voxel value represents the average Ricci-Forman curvature of all edges whose endpoints touch this voxel. Second, a voxelwise average is computed across the curvature maps of all subjects in the training set. Finally, the resulting average map is thresholded using the median of its histogram as the cutoff. This yields the “core mask” to which all subsequent analysis steps are constrained. Supplementary Figure 2 shows an example.

### Step 4 (Learning a regression model)

We use an ensemble learning approach in which multiple learners are trained independently using different subsets of the feature space [42–44]. More precisely, a large number of subsets of the feature space are randomly selected, and each subset is then used in a regression model to generate predictions of intelligence scores for the subjects of the test set. Finally, those predictions are averaged separately for each fold, see Supplementary Fig 3 for an illustration. In the experiments reported below, we found that 2000 subsets sufficed to reach convergence. In the following, those computations are described in more detail.

### Selection of edges from the core mask

A subset of the feature space is obtained by randomly selecting *m* edges whose endpoints are required to be within the core mask. In the experiments reported below, the number of edges in the core mask was around 250,000 from which *m* = 1000 edges were selected randomly for each subset. As we will show later, the VEGA algorithm is quite robust with regard to the setting of the hyperparameter *m* (Supplementary Figures 12,13,14).

### Partial least squares regression

Based on the *m* edges selected in the previous step, a linear model of the form

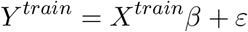

is set up to estimate the intelligence scores of the training set and learn the parameter *β*.

Here *X*^*train*^ is an *n* × *m* matrix of predictors, *Y* ^*train*^ is a vector of length *n* of intelligence scores and *ε* an error term. The entries of *X*^*train*^ are correlation values extracted from *C*_*s*_, *s* = 1, …, *n* where *n* = 325 is the number of subjects in the training set and *m* the number of randomly selected edges.

Several types of linear models might be considered for this purpose, e.g. ordinary least squares regression (OLS) or its extensions such as Lasso, Ridge, and ElasticNet. Another option would be support vector regression [45]. Here we decided to use partial least squares regression (PLS) [46, 47] because it is particularly well suited to handle problems where the predictors are highly collinear and where number of independent variables greatly exceeds the number of data points [48]. As noted above, we typically select *m* = 1000 edges, while the number of subjects in the training set is only *n* = 325 so that *m* ≫ *n*. PLS projects both *X* and *Y* to a new space that maximizes their covariance so that the predictors are projected into directions that make them more relevant for the prediction.

The PLS model is defined as

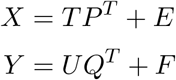

where *T, U* are projections of *X* and *Y* whose covariances are maximized. *P, Q* are their loading matrices, and the matrices *E, F* are error terms. The dimension of matrix *P* is *m* × *p* where *p* represents an intrinsic dimensionality. In the experiments reported below, we used *p* = 10. However, the results were quite robust with respect to the choice of this parameter (Supplementary Figures 12,13,14). The matrix *Q* is of dimension *m* × *q* where *q* is the number of columns in *Y*. In the present case, *Y* is a vector containing intelligence scores so that *q* = 1.

There exist several methods for computing PLS. Here we implemented the SIMPLS algorithm [48]. SIMPLS iteratively computes projections *T, U* and loading matrices *P, Q* together with a weight matrix *R* such that *β* = *RQ*^*T*^ and hence *Y* ^*train*^ ≈ *X*^*train*^ *β*.

Prior to PLS regression, the matrices *X*^*train*^ and *X*^*test*^ are row and column-centered. For details on this point, see equations 2,3,4 on the last page of the supplementary material. Likewise, the vector *Y* ^*train*^ is mean-centered and its mean is used to center the test vector *Y* ^*test*^.

#### Apply learned parameters to the test set

Once *β* has been learned from the training data, predictions for the test subjects can be obtained using *Y* ^*test*^ ≈ *X*^*test*^ *β*. Finally, the predictions are averaged within each fold.

### Benchmarks

We implemented a range of methods for benchmarking using the scikit-learn library [52, 53]. All methods are derived from the general framework called “connectome-based predictive modeling (CPM)” [3, 5, 6, 54]. CPM reflects the state of the art in the field of predicting behaviour from fMRI data and can therefore serve as a useful point of reference.

The framework consists of the following processing steps. First, a dimensionality reduction is performed by some form of brain parcellation or ICA decomposition [55, 56]. Second, a whole-brain connectivity pattern is calculated by correlating the fMRI activity time courses of every pair of parcels or components extracted in the first step. Third, a linear model relates this pattern to the behavioral score of interest. Finally, the model is applied to previously unseen data to generate a behavioral prediction.

The methods included into this benchmark differ with regard to the input data representation. The first set of benchmark methods expects input data represented in the voxel space. For these methods, we include two types of parcellations, namely the parcellations by Shen et al [49], and that by Power et al. [50].

The second set of benchmark methods expects input data to be represented in a surface representation called “ grayordinate space (CIFTI)” which is specific to the Human Connectome Project (HCP). In grayordinate space, two types of dimensionality reduction are most common. The first is the atlas-based multimodal parcellation (MMP) [51]. The second is derived from ICA decompositions so that a node is defined as an ICA component. ICA decompositions were included in our benchmark because they were used in the “megatrawl” release [2], and are thus a highly relevant reference in our context. However, analogous ICA decompositions are not available for task fMRI data in the HCP database and were thus only included as a benchmark for resting state data.

Furthermore, we included three measures of connectivity. Those are partial correlation, Pearson linear correlation and tangent correlation [57]. For all three measures, covariances are estimated using the Ledoit Wolf method [58]. In grayordinate space, we only used the tangent correlation as this was expected to produce the best results [14].

To establish the linear model, we used ridge regression throughout and relied on a generalized crossvalidation procedure over the training set to select the regularisation parameter [59]. This method provided the best results over appropriate CPM alternatives and reinforced earlier observations about its performance benefits [14]. Therefore, no feature selection step was performed as in [5].

For the benchmark we used the same fMRI input data as for the VEGA algorithm. The pre-processing was done as follows. Detrending and a high-pass filter with a cutoff frequency of 1/100 Hz was applied to remove baseline drifts. Images were standardized (zero mean with unit variance). For the voxel-space methods, a spatial Gaussian smoothing using fwhm=6mm was applied. To extract parcellated time-series, preprocessed images were signal averaged within parcels defined by either the 360 parcels of the MMP atlas [51], the 268 parcels atlas of [49] or the 264 parcel atlas of [50]. In the latter case, parcels were defined as spheres with 5mm radii around the coordinates specified in [50]. The brain atlases of [49, 50] are defined at a resolution of (2*mm*)^3^ so that no downsampling of the fMRI data in the voxel-space was applied.

Table 2 gives an overview of all methods included into the benchmark.

**Table 2:**
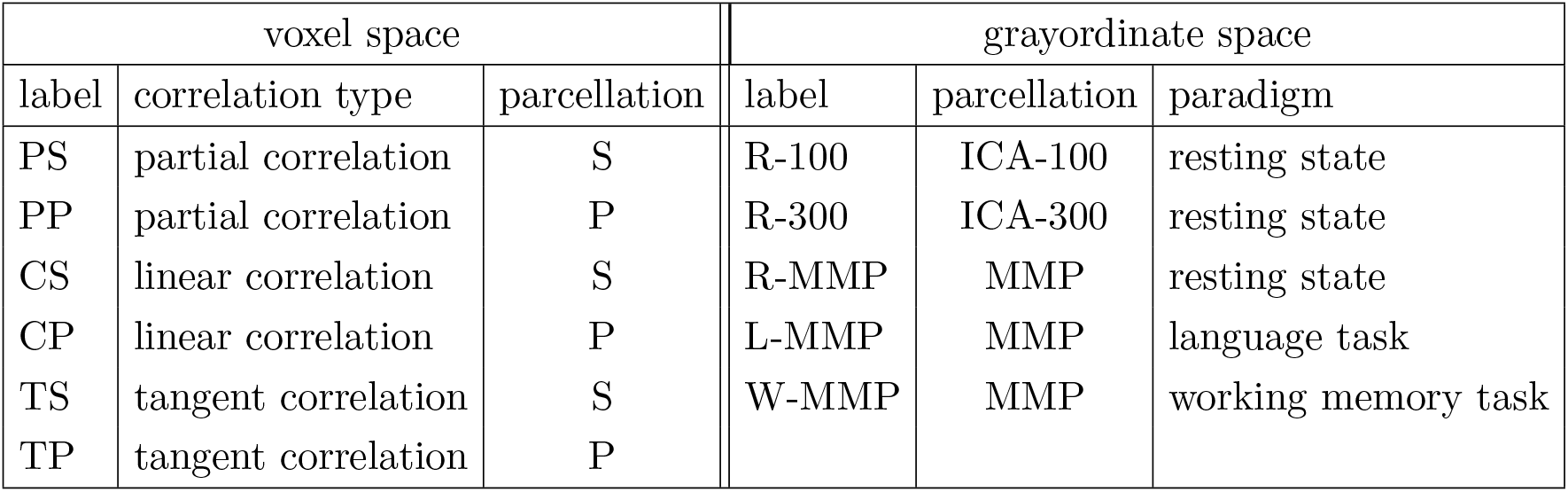
Baseline methods included into the benchmark. Our proposed algorithm VEGA was tested against a battery of competing methods that are based on the CPM framework using various connectivity types and parcellation schemes as described in the text. The parcellations are denoted as S [49], P [50] and MMP [51]. For resting state data in grayordinate space, we also used ICA decompositions with 100 and 300 nodes (ICA-100, ICA-300). The correlation types are partial correlation, Pearson linear correlation and tangent correlation [14]. In grayordinate space, tangent correlation was used throughout. The labels correspond to those in Figures 3,6,5,4.

| voxel space |  |  | grayordinate space |  |  |
| --- | --- | --- | --- | --- | --- |
| label | correlation type | parcellation | label | parcellation | paradigm |
| PS | partial correlation | S | R-100 | ICA-100 | resting state |
| PP | partial correlation | P | R-300 | ICA-300 | resting state |
| CS | linear correlation | S | R-MMP | MMP | resting state |
| CP | linear correlation | P | L-MMP | MMP | language task |
| TS | tangent correlation | S | W-MMP | MMP | working memory task |
| TP | tangent correlation | P |  |  |  |

## Results

In our experiments, we used 390 subjects and 6-fold cross validations so that each training set had 325 subjects, and each test set had 65 subjects. We recorded the Pearson linear correlation and the predictive coefficient *R*^2^ between the observed and the predicted intelligence scores where *R*^2^ = 1 − *SS*_*res*_*/SS*_*total*_. Here *SS*_*res*_ is the residual sum of squares, and *SS*_*total*_ is total sum of squares. We also report the mean absolute error (MAE). We randomly defined 20 different training/test splits resulting in 6 × 20 = 120 different correlation scores, *R*^2^ and MAE values. We used the same training/test splits for the benchmark methods described in the previous section.

Figures 3,4,5,6 show the resulting Pearson correlations between the observed and the predicted intelligence scores for the G-factor, total, crystallized and fluid intelligence, respectively. The corresponding *R*^2^ values and mean absolute errors are in the supplementary material (Supplementary Figures 4,5,6,7). We first note that the results obtained by VEGA in the two task conditions were clearly better than the results in the resting state condition.

**Figure 3:**
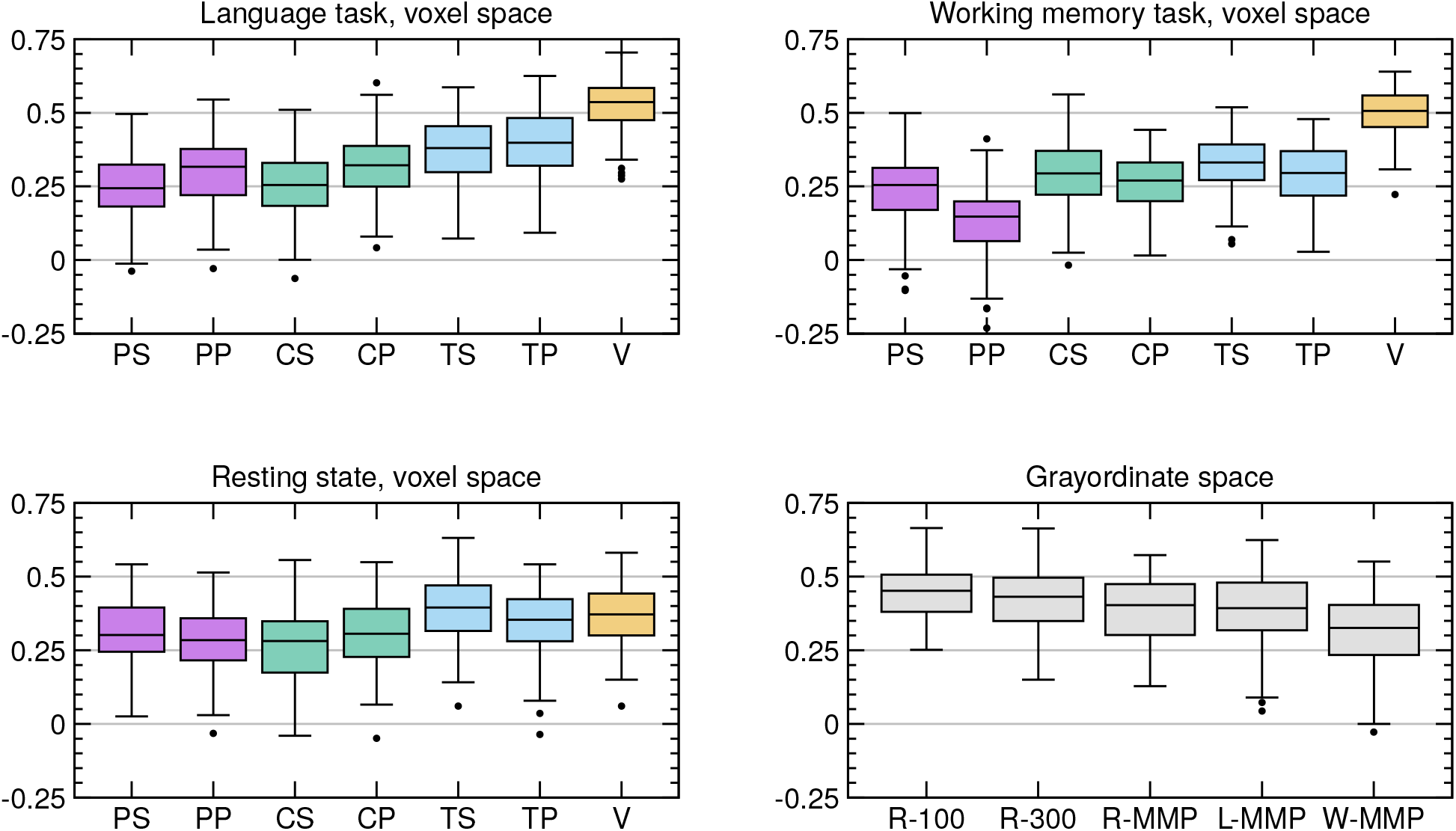
Correlations between observed and predicted G-factor. The boxplots show the Pearson linear correlations between the observed versus predicted intelligence scores of 65 test subjects resulting from 6-fold crossvalidations in 20 different train/test splits (6 × 20 = 120 correlation values). The corresponding R^2^ values are in the supplement. The results of the new method VEGA (‘V’) are shown in orange. It was tested against several competing methods using the same data and train/test splits, see table 2. Note that in the language and working memory tasks, VEGA outperformed all competing methods. In resting state data (voxel space), its accuracy is comparable to the best competing methods.

**Figure 4:**
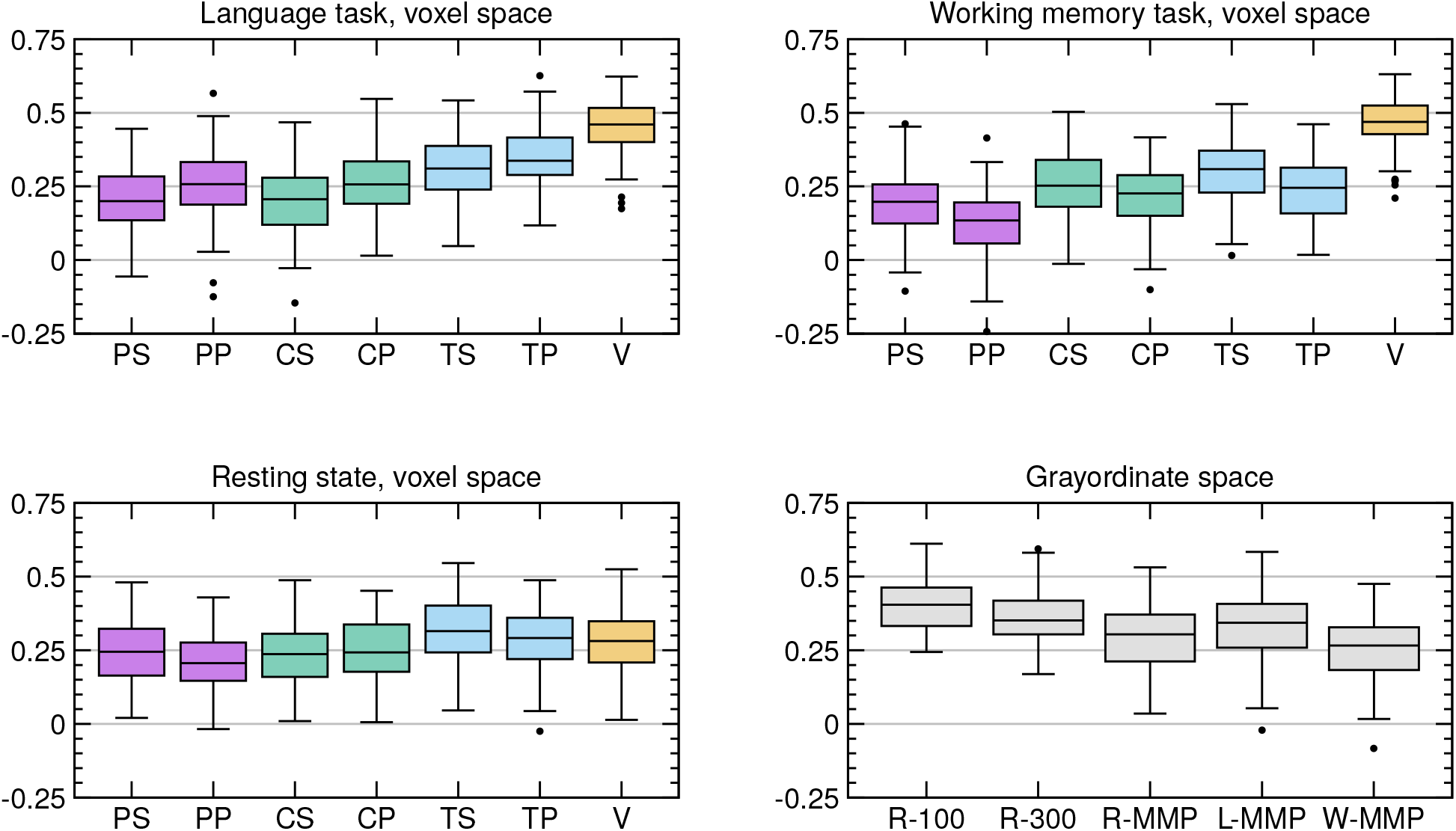
Correlations between observed and predicted CogTotal. Results for the CogTotal score, see caption of Figure 3 for more details.

**Figure 5:**
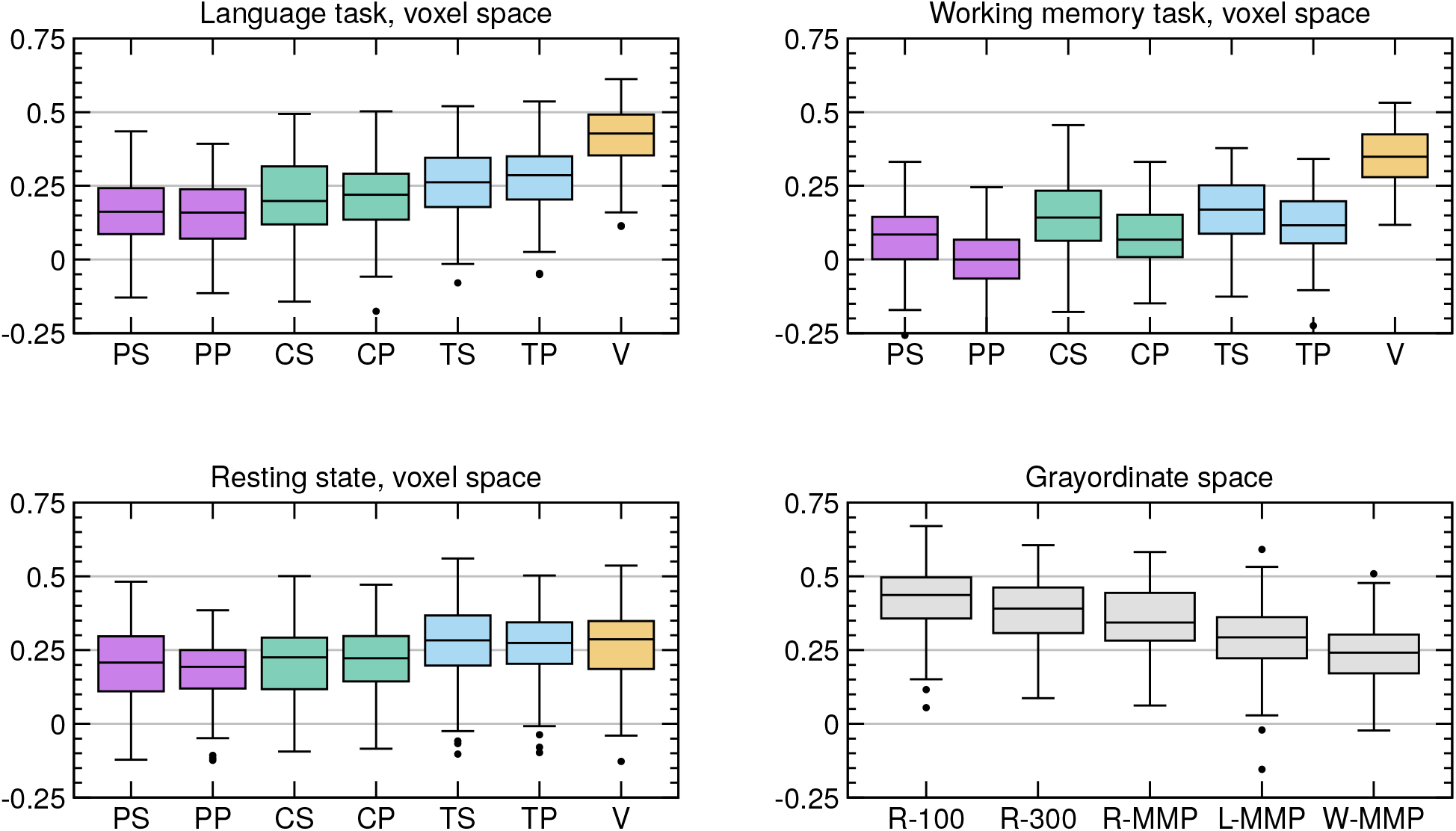
Correlations between observed and predicted CogCrystal. Results for the CogCrystal score, see caption of Figure 3 for more details.

**Figure 6:**
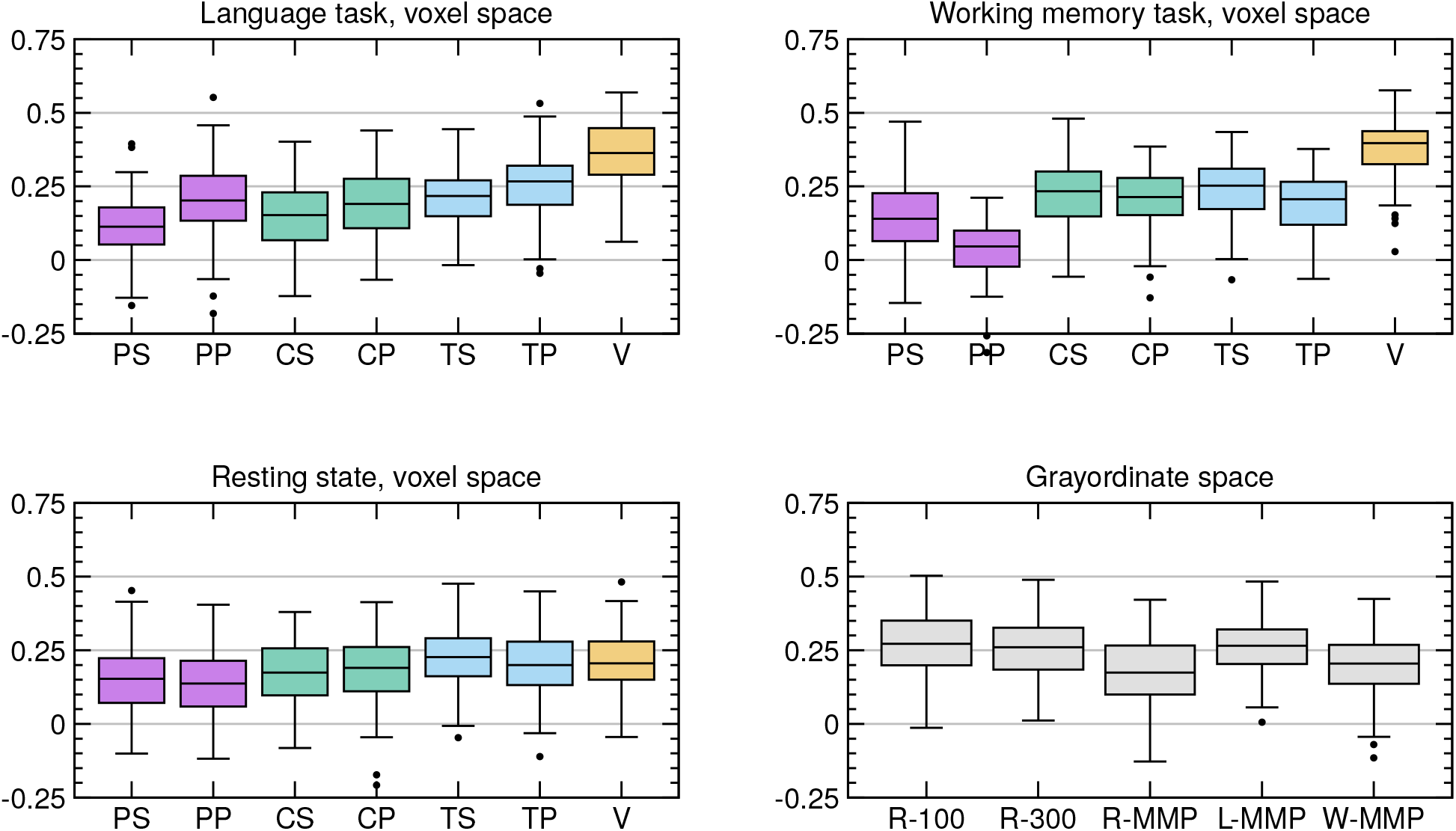
Correlations between observed and predicted CogFluid. Results for the CogFluid score, see caption of Figure 3 for more details.

All results were obtained with *m* = 1000, *p* = 10 as hyperparameters where *m* denotes the number of edges randomly selected in each subset of the ensemble learning, and *p* denotes the intrinsic dimensionality of the PLS regression. We then investigated the influence of those two hyperparameters. We found that the results remained almost unchanged with *m* = 100, 500, 1000, 2000, 5000, and *p* = 3, 10, 20, 50. At *m* = 100 the prediction accuracy decreased, see Supplementary Figures 12,13,14.

Figure 7 shows scatter plots from one of the twenty training/test splits obtained by VEGA. Here we also included an average of the results between the two tasks. A systematic evaluation of the effect of combining the two tasks is shown in Supplementary Figure 15. For example, the median correlation between observed and predicted G-factor improved to 0.587 (*R*^2^=0.294). And the median correlation between the observed and predicted CogTotal improved to 0.526 (*R*^2^=0.241).

**Figure 7:**
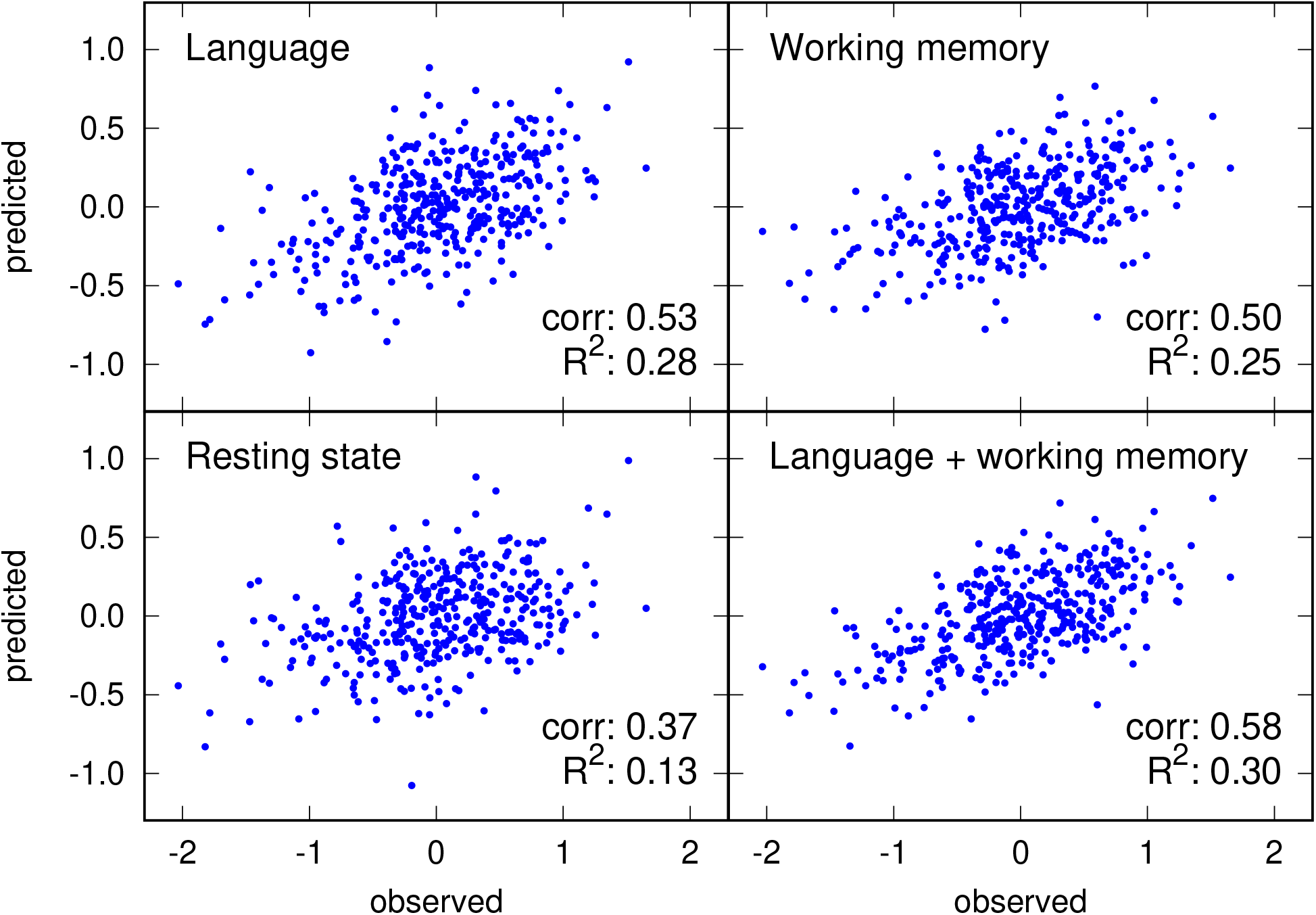
Scatter plots showing predictions of the G-factor. Scatter plots for the prediction of the G-factor obtained by the proposed algorithm VEGA are shown. Each dot represents one of the 390 test subjects. Note that the results from the two tasks are markedly better than that of the resting state. The average of the two tasks yields the best result. All four scatter plots are derived from the same training/test split.

The boxplots of Figures 3,4,5,6 show that VEGA outperformed all benchmark methods in the two task conditions. To assess statistical significance, we additionally performed an inference for the generalization error based on the Z-transformed Pearson correlation scores. Note that standard t-tests are not valid in this context because the results are derived from the same pool of subjects so that independence assumptions are violated. Therefore, we used a modified t-test that corrects for this problem [60, 61]. In comparing the two task-based results of VEGA against the best voxel-based benchmarks, we found that VEGA was indeed significantly better in all cases (*p* < 0.025). We then combined the two task-based results of VEGA and compared them against all benchmark methods including the ones obtained in the grayordinate space, and found that the VEGA result was significantly better throughout (*p* < 0.015).

Furthermore, we investigated the effect of scan time on prediction accuracy in the resting state condition. For example, we found that the median correlation between observed and predicted G-factor declined slightly from 0.371 (full scan time, 58 min) to 0.360 and 0.347 when the scan time was cut in half (29 min for each session). And it declined to 0.286, 0.306, 0.294, 0.258 when the scan time was reduced even further (14.4 min for each run). For details, see Supplementary Figure 16.

We then tested whether or not Ricci-Forman curvature actually helped to increase prediction accuracy. In the language task, we found that without Ricci-Forman curvature the results for the prediction of the G-factor (median correlations and median *R*2) decreased from 0.54 (0.27) to 0.50 (0.23). In the working memory task, it decreased from 0.51 (0.23) to 0.49 (0.20). For rsfMRI, it did not produce a better prediction. For details, see Supplementary Figure 17.

Finally, we investigated which brain areas are most relevant for predicting intelligence. Figure 8 shows these areas for the G-factor in the language task. Specifically, we compute the mean factor loadings of *X* that are recorded in the load matrix *P*. We distinguish between positive and negative factor loadings so that for each edge *i*, we have

**Figure 8:**
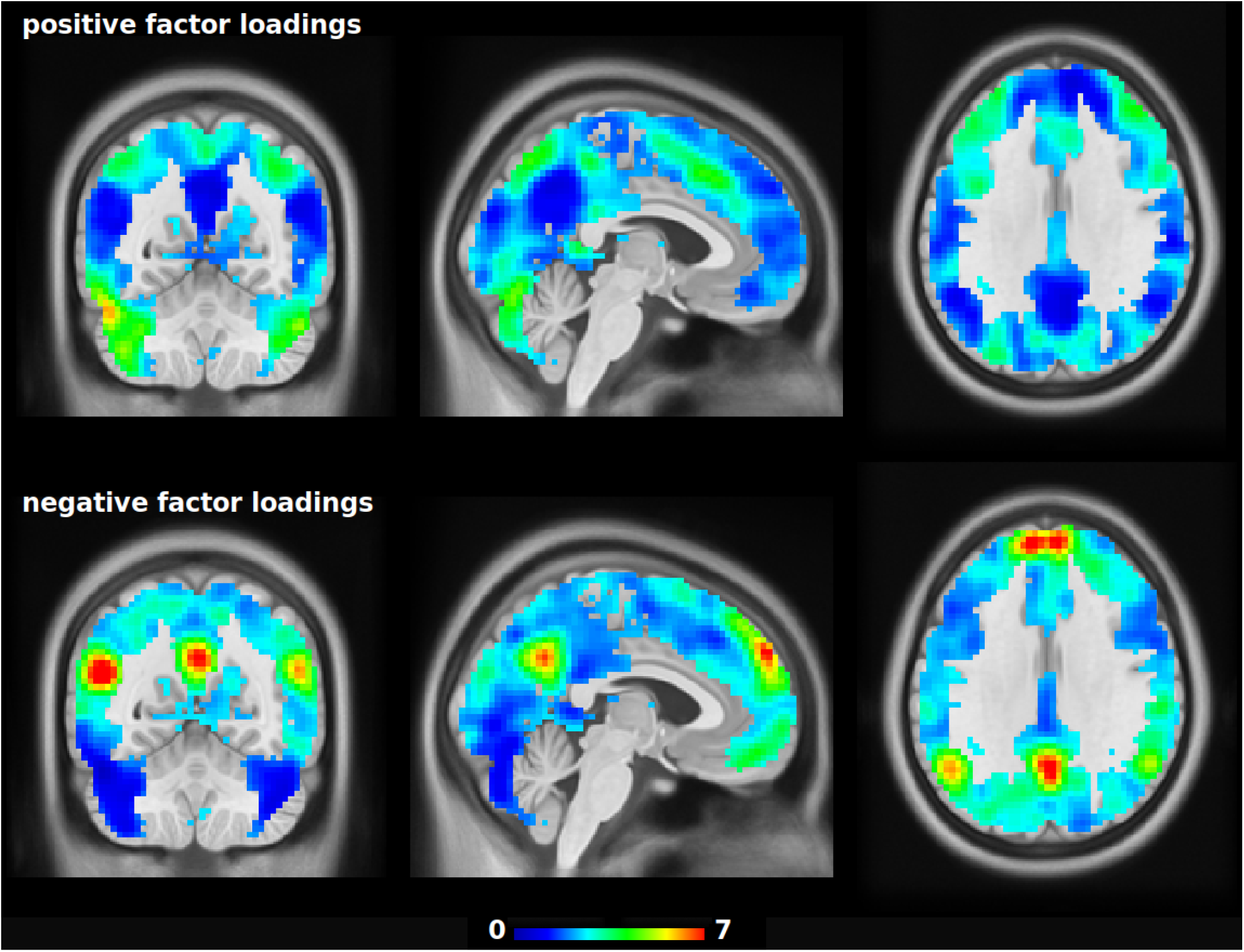
Predictive areas for general intelligence in the language task. The colors encode factor loadings (matrix P) estimated by partial least squares regression averaged over all folds. Strong positive loadings indicate areas where connectivity with other brain regions is positively correlated with general intelligence. Strong negative loadings indicate areas where connectivity with other brain regions is negatively correlated with general intelligence. For example, the negative loadings seem to highlight the default mode network (DMN). This suggests that a well connected DMN correlates with low general intelligence. A spatial Gaussian filter (fwhm=7mm) was applied for better visualization. Note that the colors only show relative weights, they do not have interpretable units.

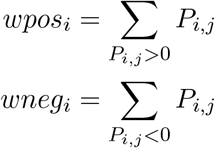

where the index *j* = 1, …*p* denotes the latent factors. The weights *wpos*_*i*_, *wneg*_*i*_ are then mapped onto a positive and a negative weight map as shown in Figure 8. Supplementary Figures 18, 19 show similar images for the working memory task and the resting state data.

## Discussion

We have introduced a new machine learning method for predicting intelligence from fMRI data. In contrast to existing methods, it does not require a presegmentation or a brain atlas, nor does it depend on an ICA decomposition. Rather, it handles the high dimensionality and multicollinearity of the data via partial least squares regression and ensemble learning.

Recently, Faskowitz et al. [62] have noted that the traditional metric of functional connectivity should be complemented by more complex edge weights and introduced an edge-centric measure. Here we have proposed Ricci-Forman-curvature as an alternative concept. It incorporates information from adjacent edges and helps to improve predictability by reducing dimensionality.

We implemented a range of existing methods for establishing a benchmark against which we compared our proposed method, see also [63]. We first note that the results in Fig 3(R-100) are approximately consistent with the results previously published by Dubois et al. [21]. Also in agreement with the literature, the benchmark showed that tangent correlation is generally superior to other measures of correlation [14]. Both observations demonstrate the replicability of earlier publications, and also the realism of our benchmark.

A comparison against the benchmark showed that the proposed algorithm VEGA offers a significant improvement in prediction accuracy in tfMRI data. This is remarkable because this improvement was achieved at a fraction of the scan time required for rsfMRI. Specifically, the scan time of the rsfMRI data was almost 1 hour, whereas the scan time for the language task was only about 7.5 minutes, and for the working memory task is was about 10 minutes. Dubois et al [21] report a noticeable decline in prediction accuracy when the scan time in rsfMRI was reduced to about 30 minutes, an observation that is supported by our own data (Supplementary Fig. 16). This suggests that rsfMRI may be too unspecific to permit good prediction accuracies at scan times that will eventually be feasible in clinical applications.

Therefore, we believe that tfMRI may be a better choice for predicting individual behaviour provided the task is suitable for the application at hand. This is in line with a new initiative for evaluating the reliability of tfMRI data [64], and it agrees with Greene et al. [65] who reported that task-induced brain state manipulation improves prediction of individual traits. Surprisingly, the methods included in our benchmark failed to exploit the advantages of tfMRI data. We speculate that dimensionality reduction via brain parcellations may be the primary reason for this failure. In essence, current parcellation schemes may not be able to reflect the continuous nature of brain topography [66].

The brain areas most relevant for intelligence prediction in the language task appear biologically plausible as success in intelligence testing is based on focused mental activity during specific tasks. Indeed, the strongest result regarding factor loadings (Figure 8) was a negative relationship between connectivity of brain areas typically associated with internally-oriented thought processes (i.e. mind wandering [67]) and intelligence which concurs with the fact that such processes are rather disruptive than helpful during cognitive tasks.

We conclude that the prediction of individual behaviour from fMRI data can be greatly improved with the help of new strategies. In particular, dimensionality reduction via ensemble learning and the use of task-based fMRI instead of rsfMRI appear to be the most important aspects in this endeavour.

## Software

The software will be made public soon.

## Acknowledgments

The project was partially funded by Reinhart Koselleck Project DFG SCHE 658/12.

Data were provided by the Human Connectome Project, WU-Minn Consortium (Principal Investigators: David Van Essen and Kamil Ugurbil; 1U54MH091657) funded by the 16 NIH Institutes and Centers that support the NIH Blueprint for Neuroscience Research; and by the McDonnell Center for Systems Neuroscience at Washington University.

## Supplementary Information

**Supplementary Figure 1:**
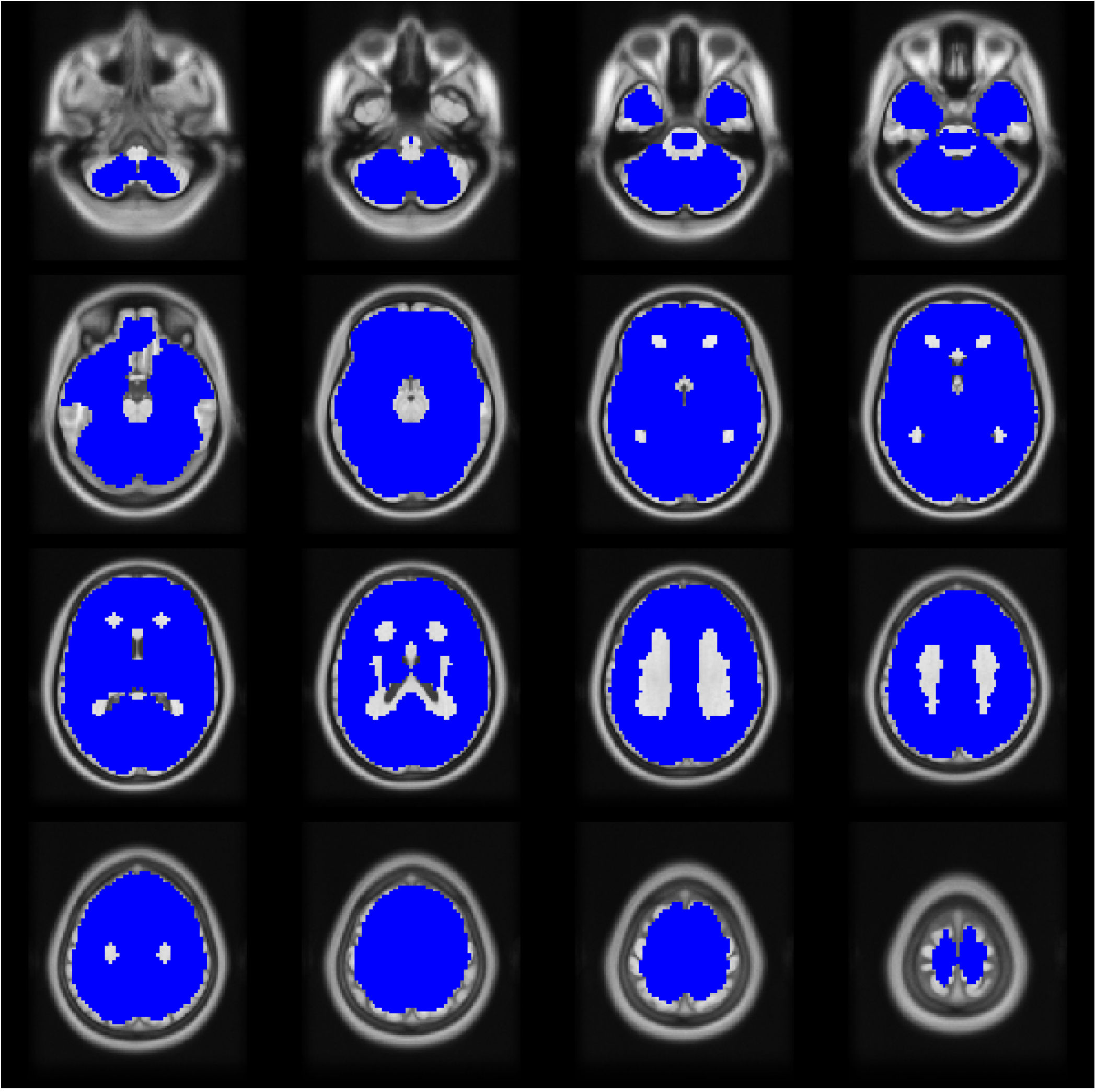
The brain mask. The brain mask used for this study. It contains 55856 voxels.

**Supplementary Figure 2:**
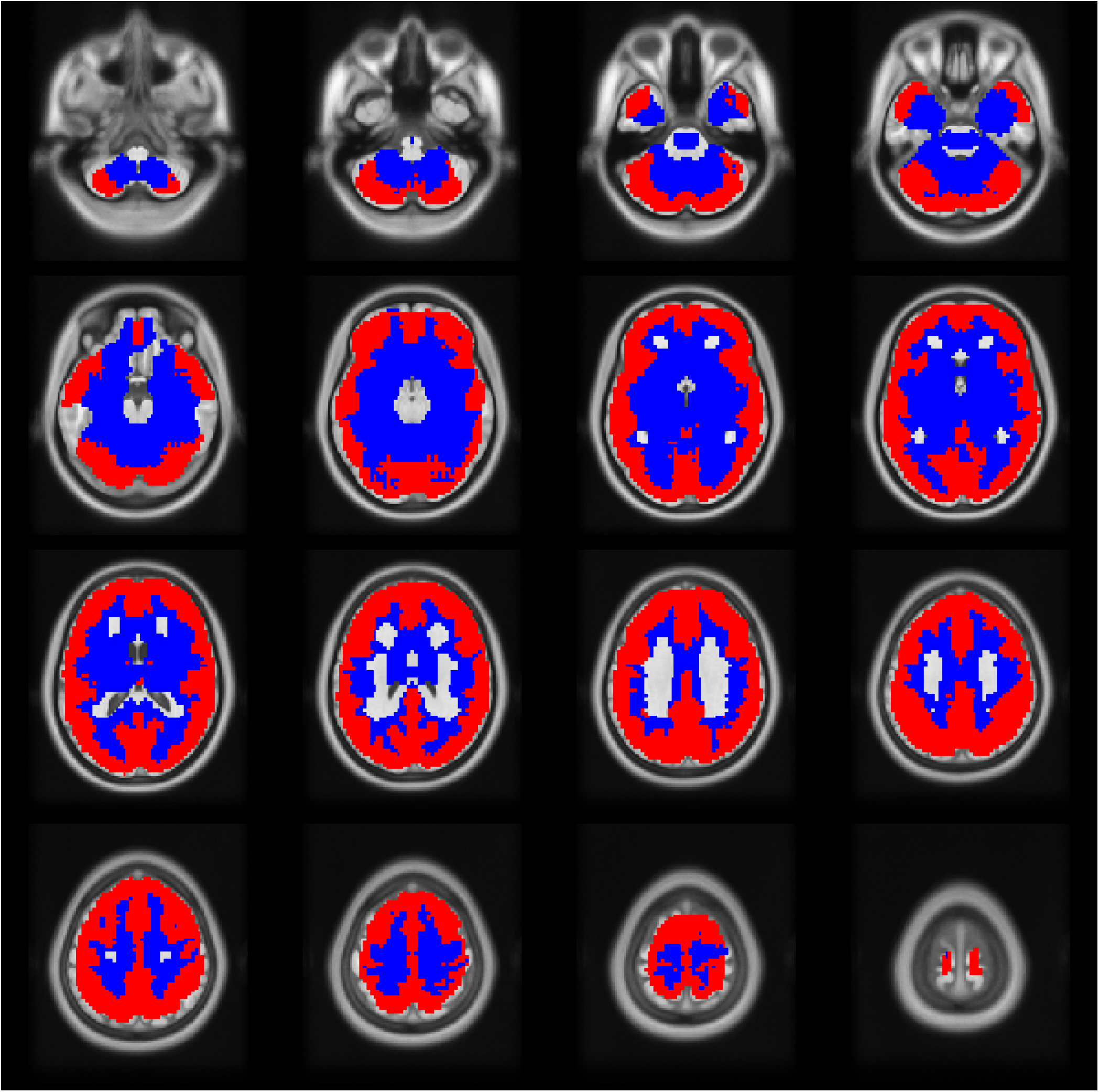
The coremap computed using Ricci-Forman curvature. The red areas show the coremap computed by thresholding the Ricci-Forman curvature map superimposed on the original brain mask shown in blue. Here, an average of the coremaps across all folds in one of the 20 train-test splits is shown. It was computed for predicting the G-factor from the fMRI data of the language task.

**Supplementary Figure 3:**
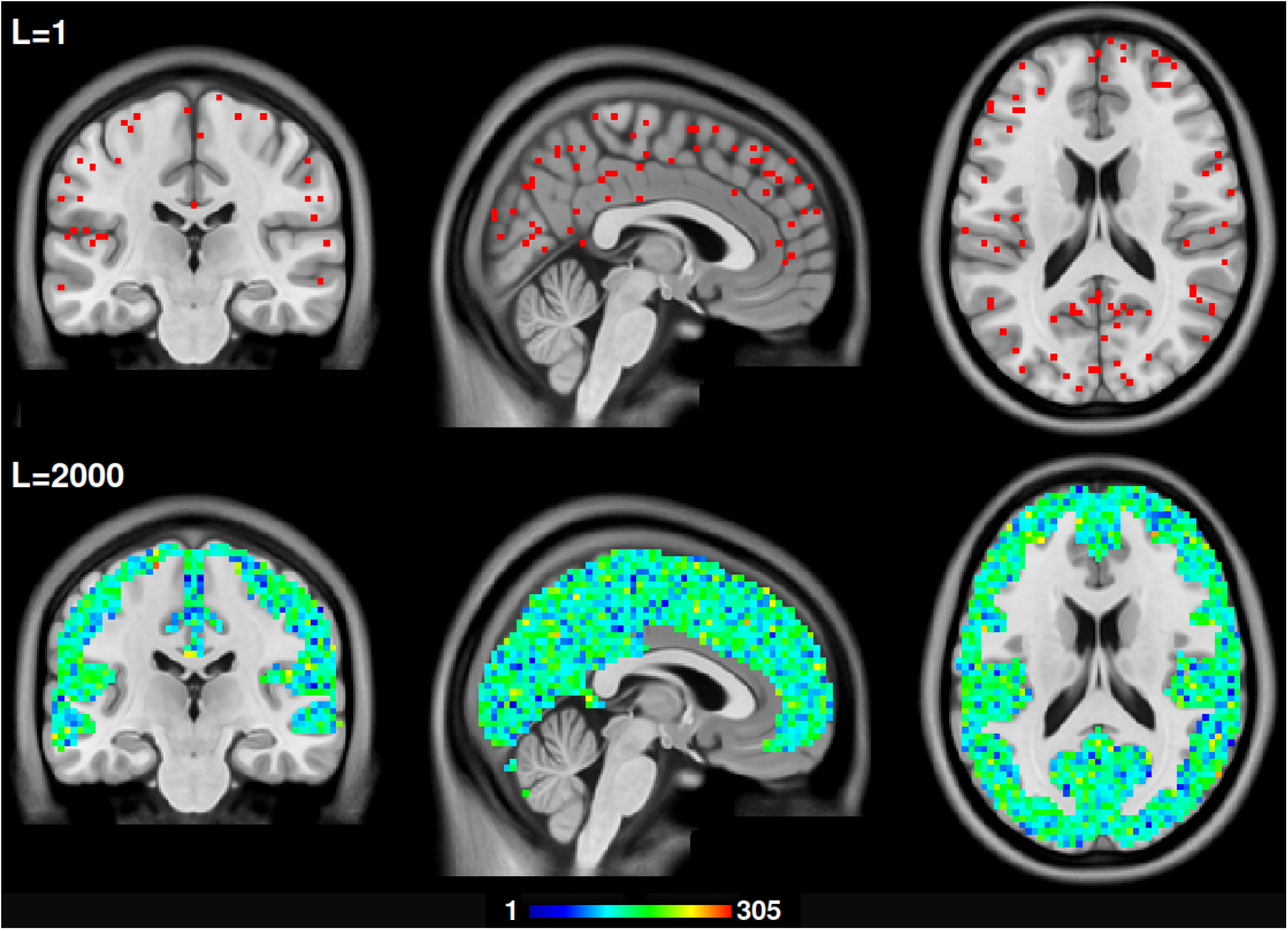
Ensemble learning. The top image shows voxels that are endpoints in one subset (L=1) of 1000 randomly selected edges used for ensemble learning. The bottom images shows the number of times voxels are visited in 2000 such subsets (L=2000). During ensemble learning, each subset is used in a regression model to derive predictions of intelligence scores. For each subject in the test set, the resulting 2000 predictions are averaged to reach a final prediction of intelligence. In this example, voxels were visited about 143 times on average (µ = 143.22, σ = 36.03). Note that the distribution of the voxels is constrained by the coremap derived from the Ricci-Forman curvature.

**Supplementary Figure 4:**
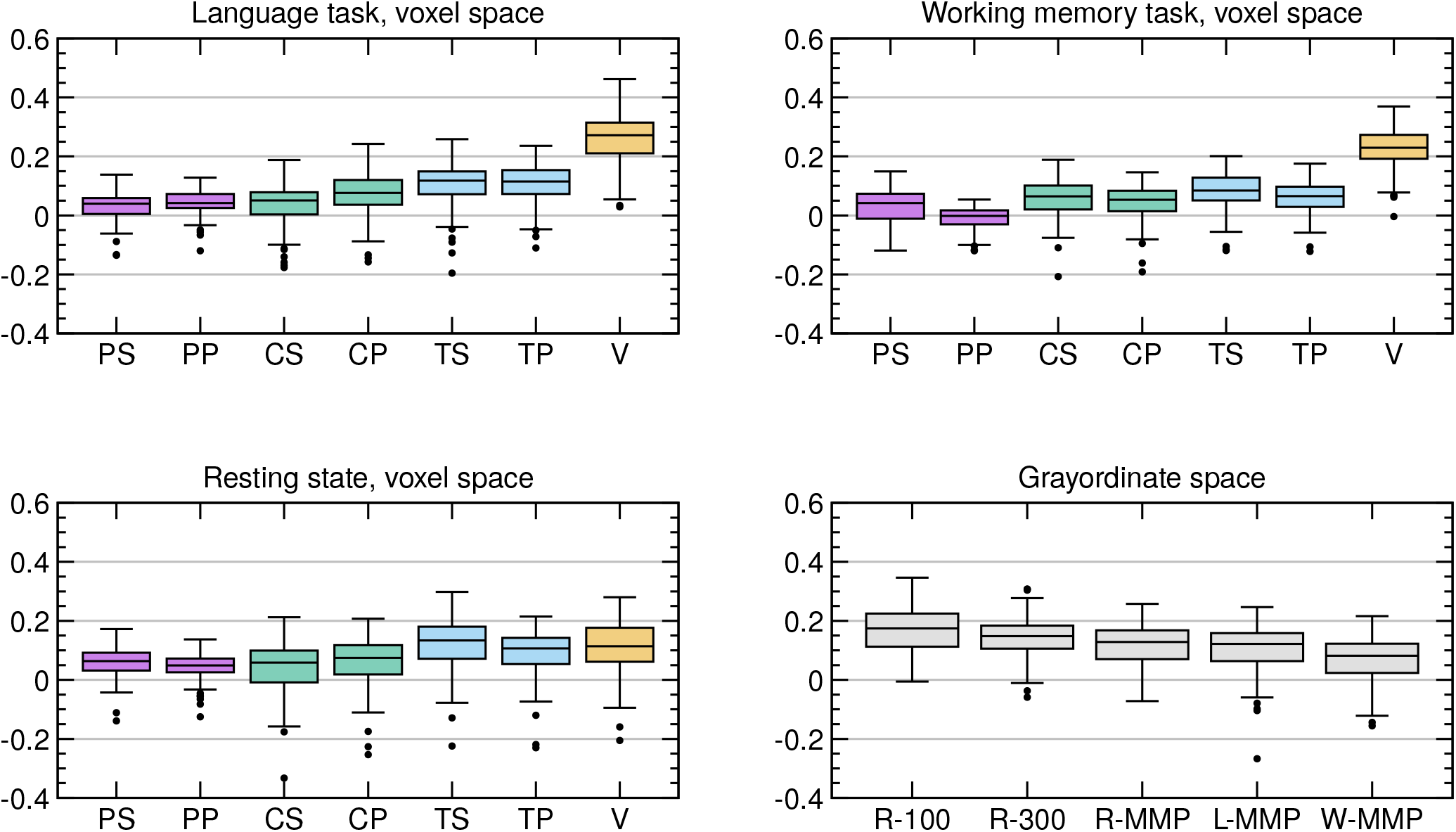
R^2^ between observed and predicted G-factor. The boxplots show the coefficient of determination R^2^ between the observed versus predicted IQ scores of 65 test subjects resulting from 6-fold crossvalidations in 20 different train/test splits (6 × 20 = 120 correlation values). The corresponding R^2^ values are in the supplement. The results of the new proposed method VEGA (‘V’) are shown in orange. It was tested against several competing methods as listed in table?? using the same data and train/test splits. Note that in the language and working memory tasks, the proposed method outperformed all competing methods. In resting state data, its accuracy is comparable to the best competing methods.

**Supplementary Figure 5:**
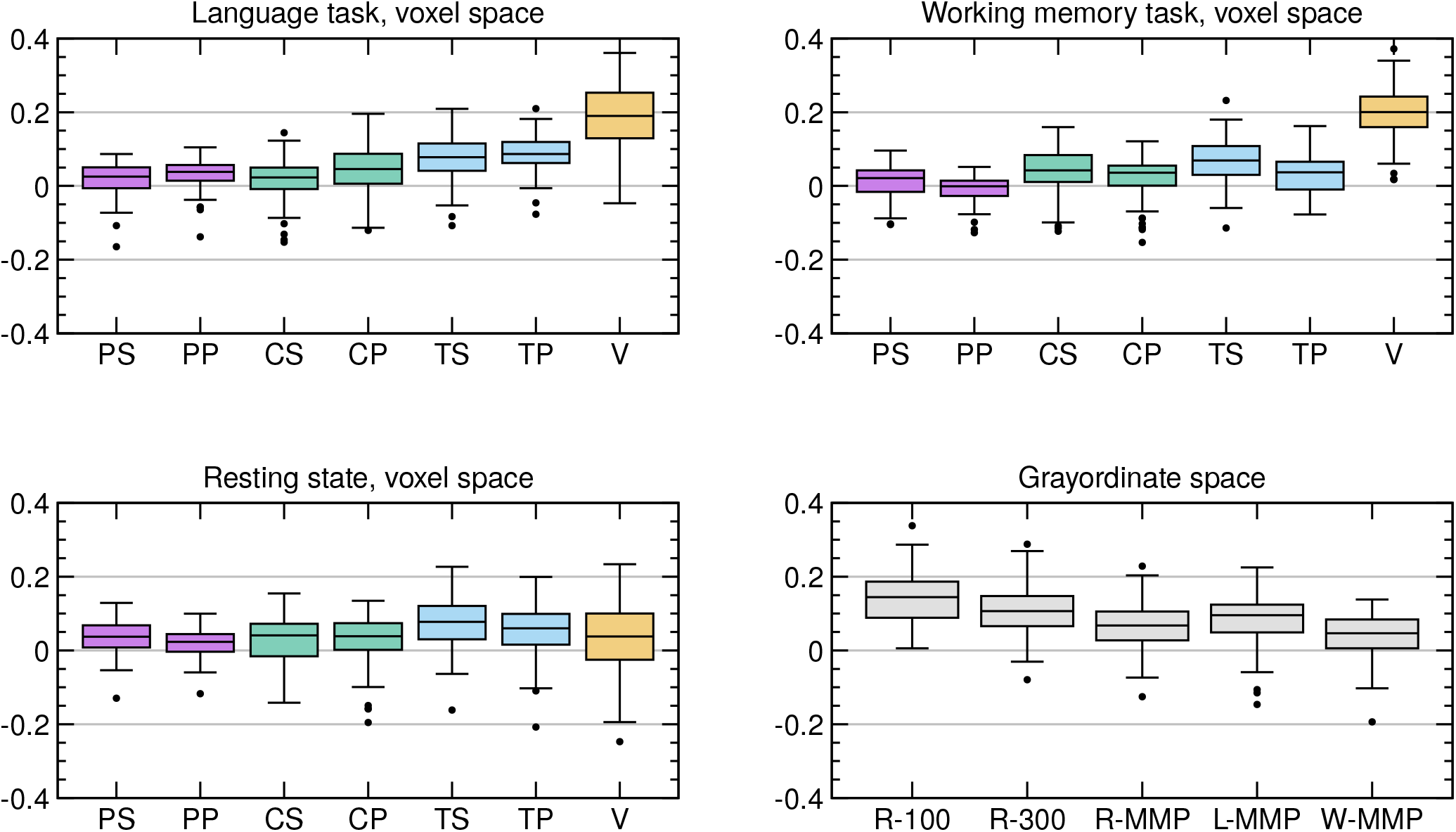
R^2^ between observed and predicted CogTotal. Results for the CogTotal score, see caption of Supplementary Figure 4 for more details.

**Supplementary Figure 6:**
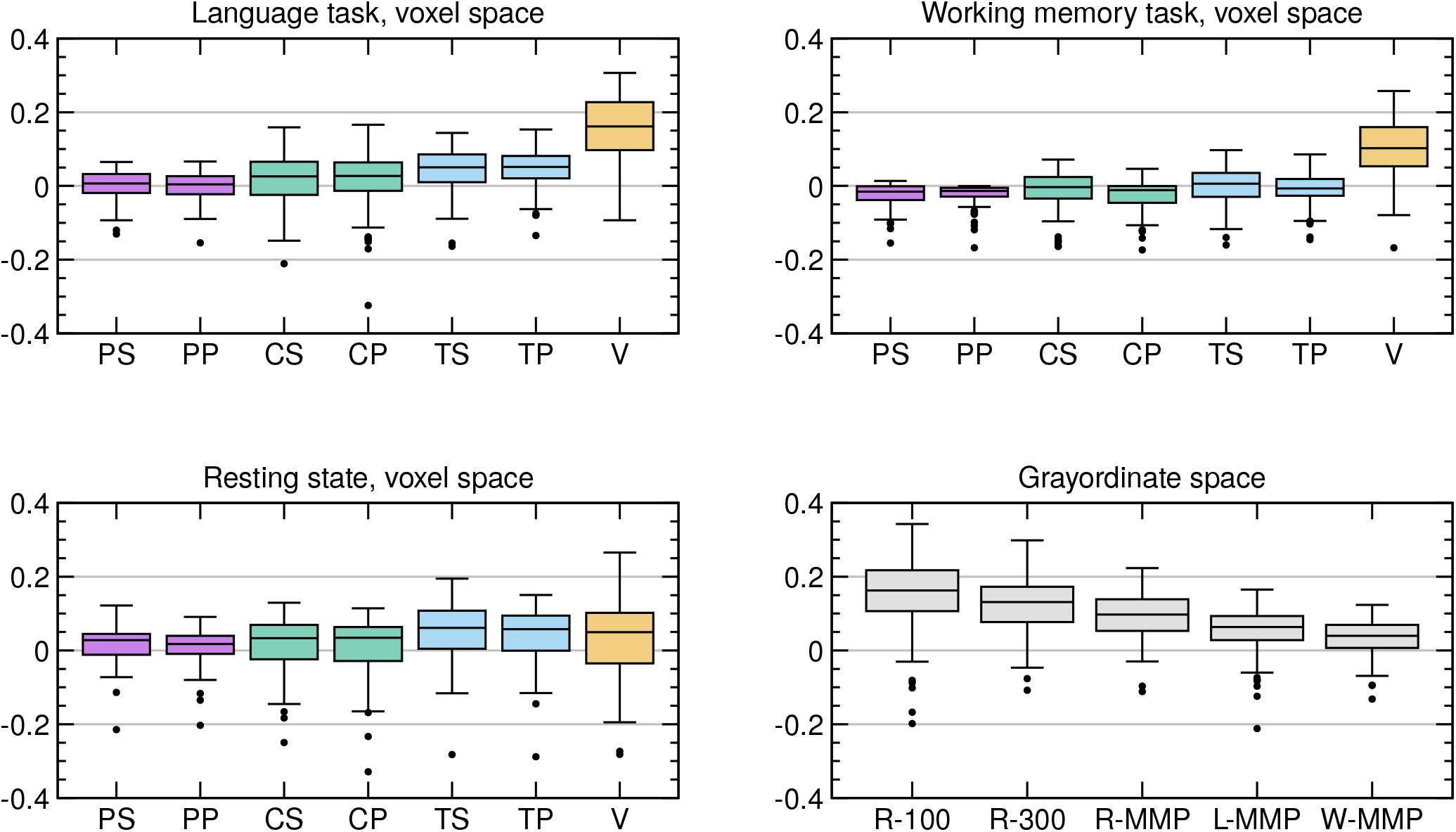
R^2^ between observed and predicted CogCrystal. Results for the CogCrystal score, see caption of Supplementary Figure 4 for more details.

**Supplementary Figure 7:**
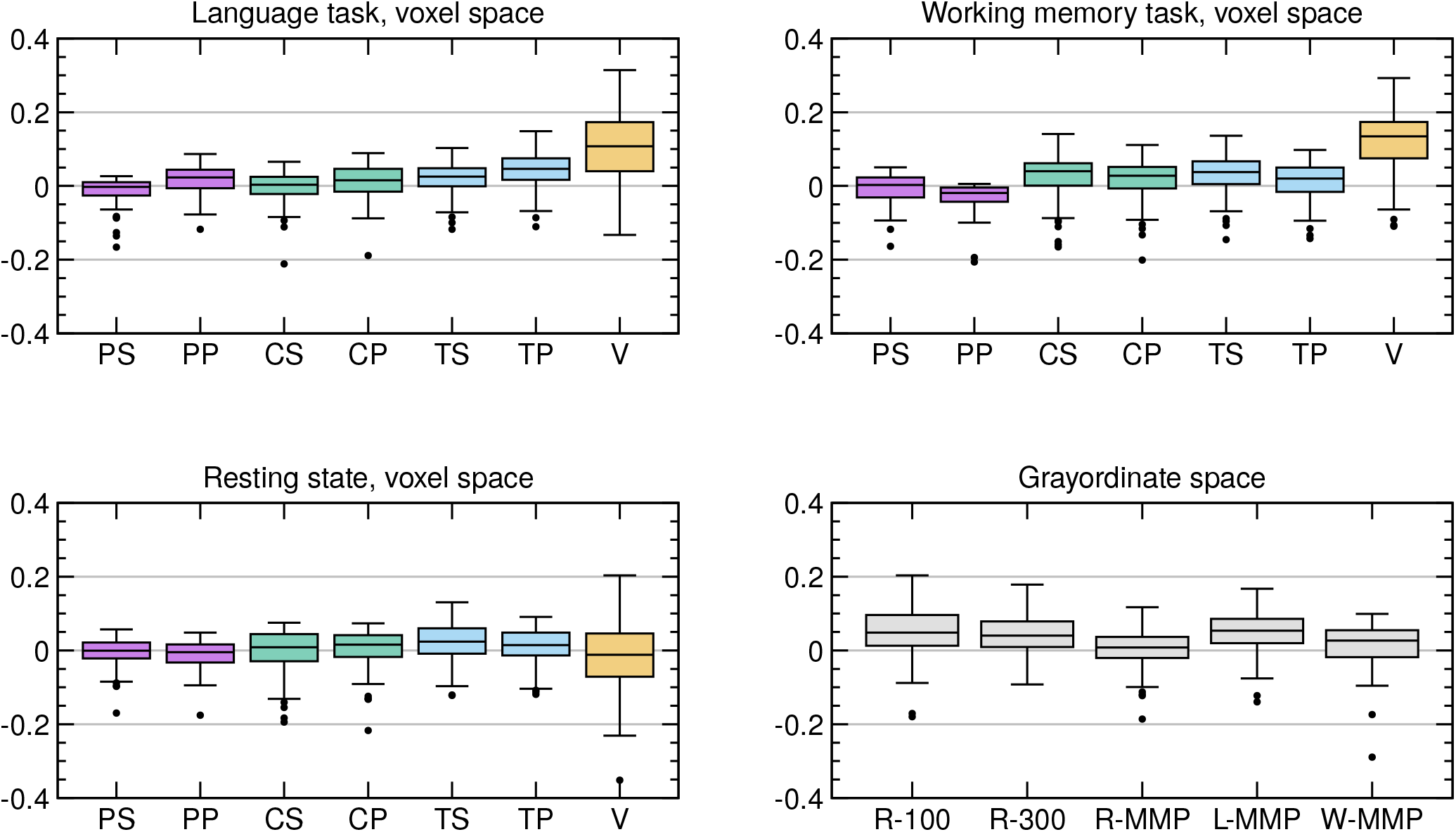
R^2^ between observed and predicted CogFluid. Results for the CogFluid score, see caption of Supplementary Figure 4 for more details.

**Supplementary Figure 8:**
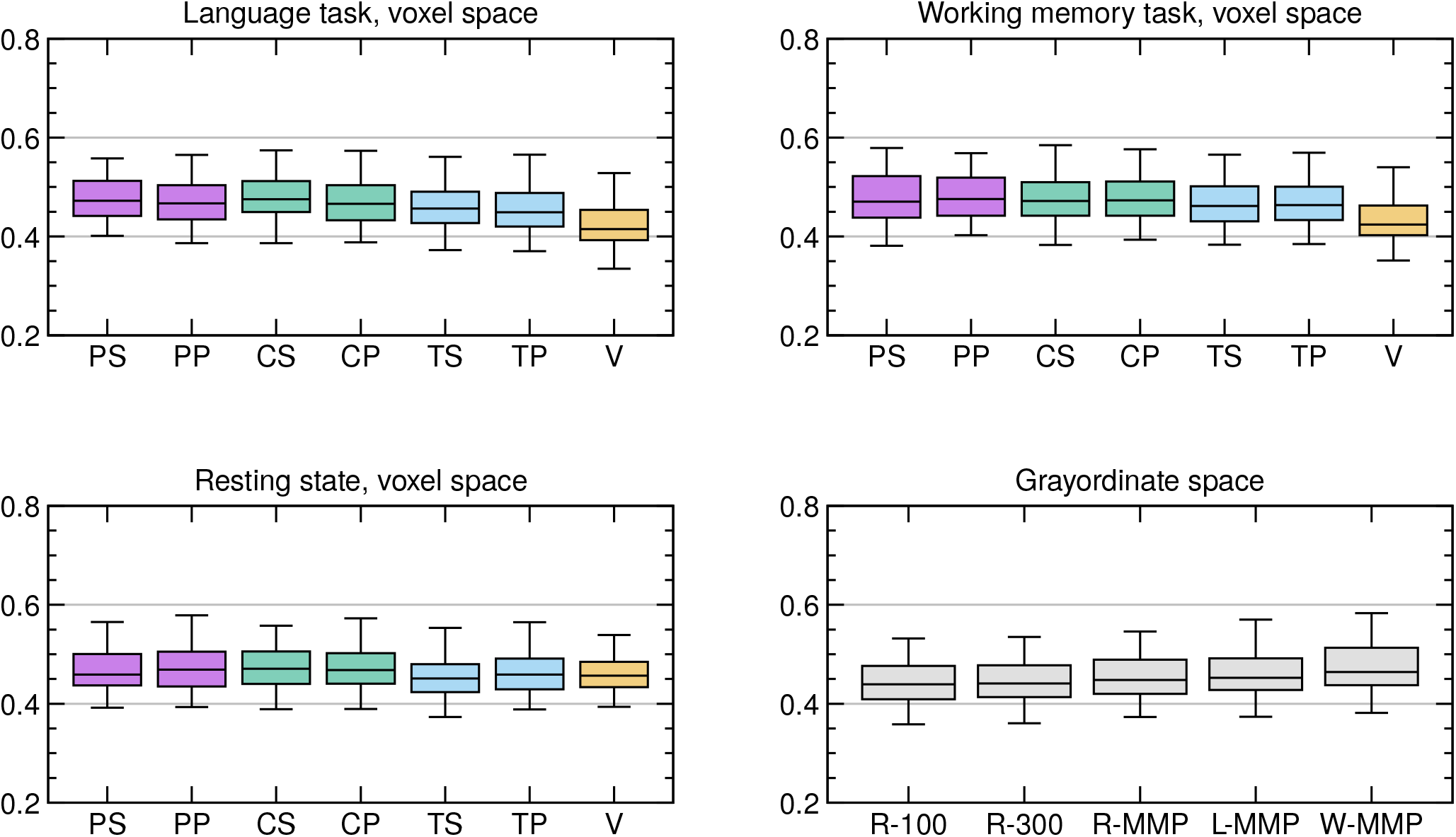
Mean absolute error (MAE) between observed and predicted G-factor. The boxplots show the mean absolute error between the observed versus predicted IQ scores of 65 test subjects resulting from 6-fold crossvalidations in 20 different train/test splits (6 × 20 = 120 correlation values). The corresponding R^2^ values are in the supplement. The results of the proposed method VEGA (‘V’) are shown in orange. It was tested against several competing methods as listed in table?? using the same data and train/test splits. Note that in the language and working memory tasks, the proposed method outperformed all competing methods. In resting state data, its accuracy is comparable to the best competing methods.

**Supplementary Figure 9:**
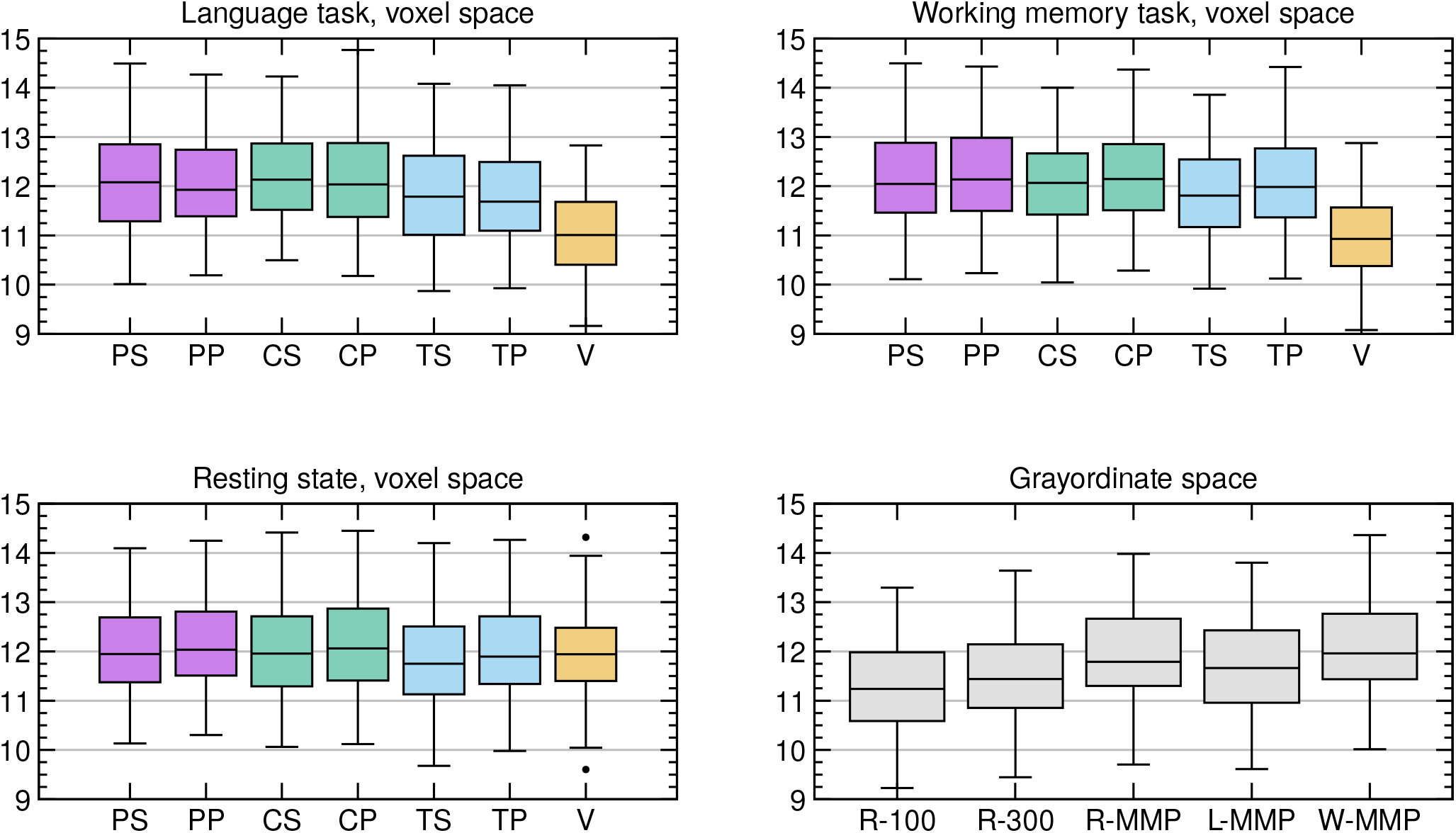
MAE between observed and predicted CogTotal. Results for the CogTotal score, see caption of Supplementary Figure 4 for more details.

**Supplementary Figure 10:**
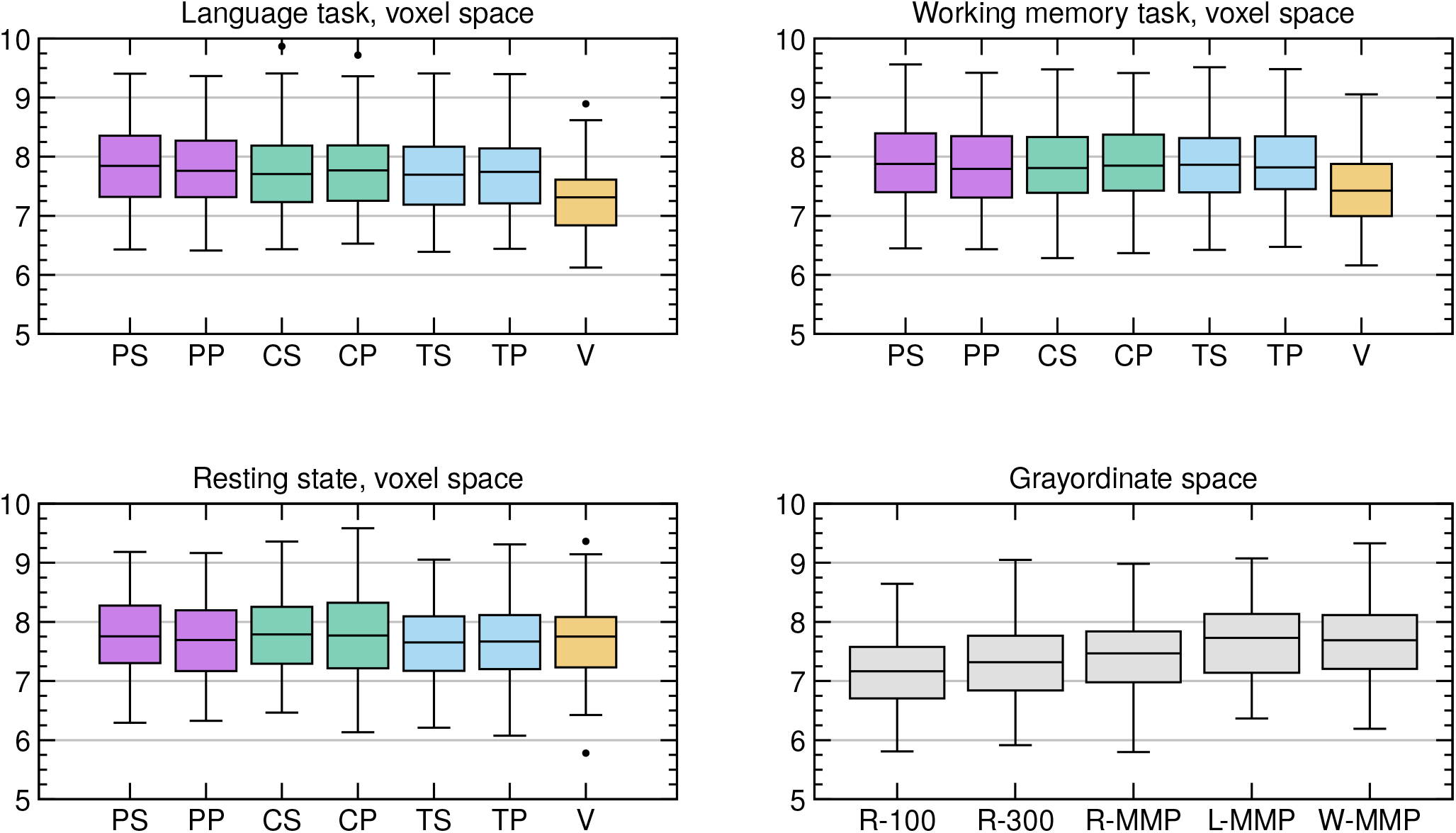
MAE between observed and predicted CogCrystal. Results for the CogCrystal score, see caption of Supplementary Figure 4 for more details.

**Supplementary Figure 11:**
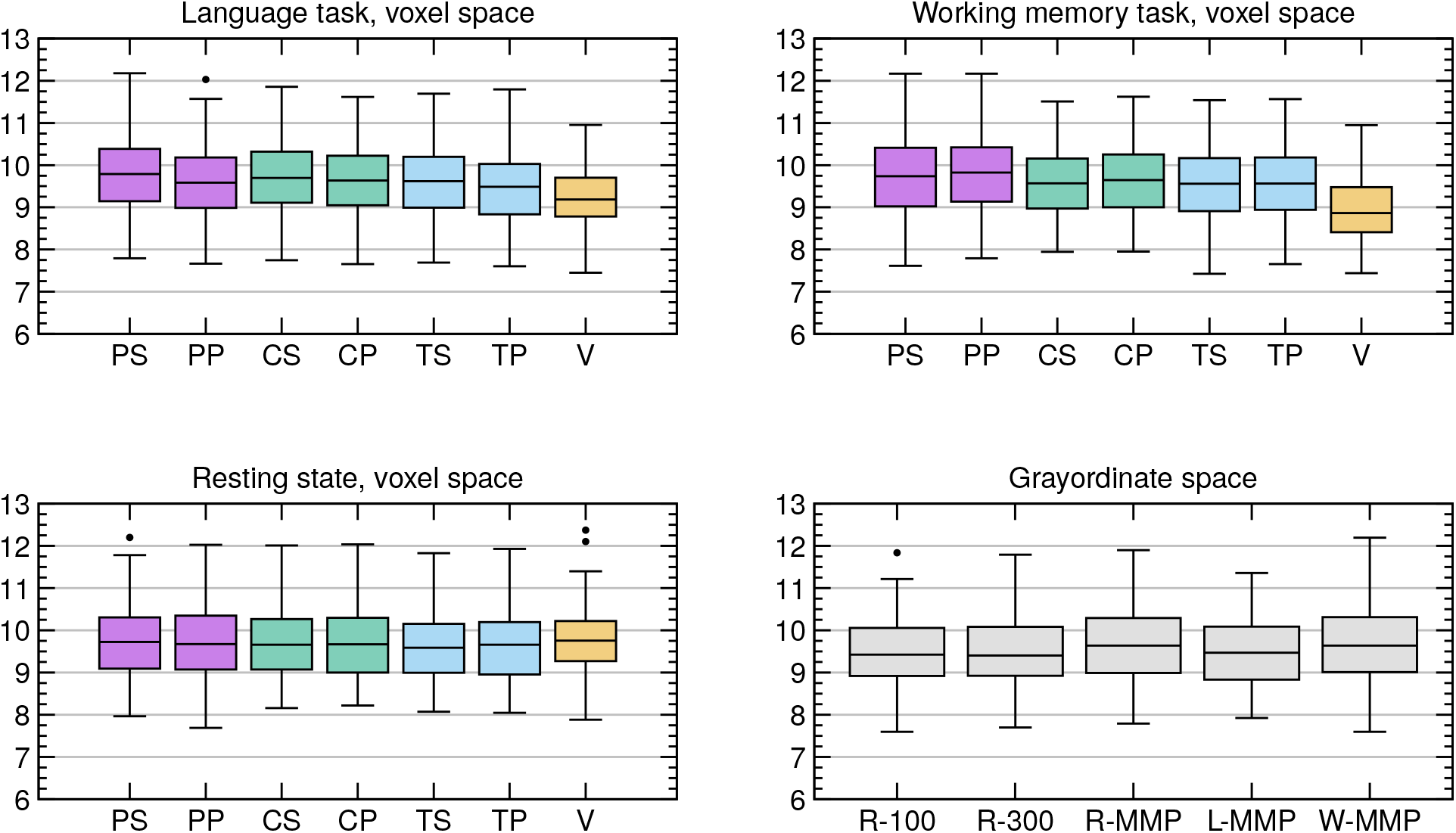
MAE between observed and predicted CogFluid. Results for the CogFluid score, see caption of Supplementary Figure 4 for more details.

**Supplementary Figure 12:**
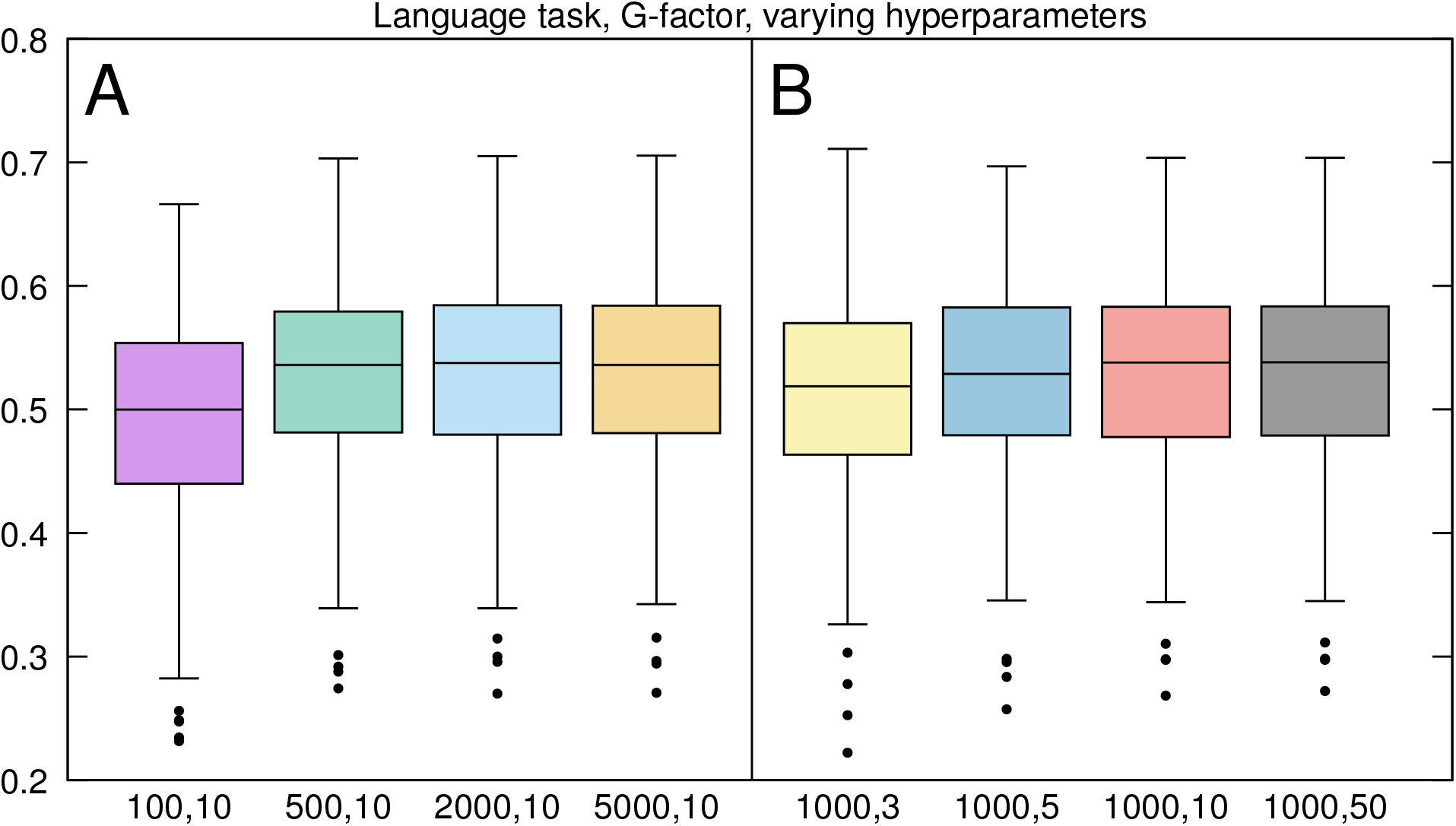
Prediction accuracy with various hyperparameter settings. The plot shows correlations between observed and predicted G-factors using the language task. The left boxplots (A) shows a variation of the parameter m = 100, 500, 2000, 5000, with p = 10 fixed. The right plot (B) shows a variation of the parameter p = 3, 5, 10, 50 with m = 1000 fixed.

**Supplementary Figure 13:**
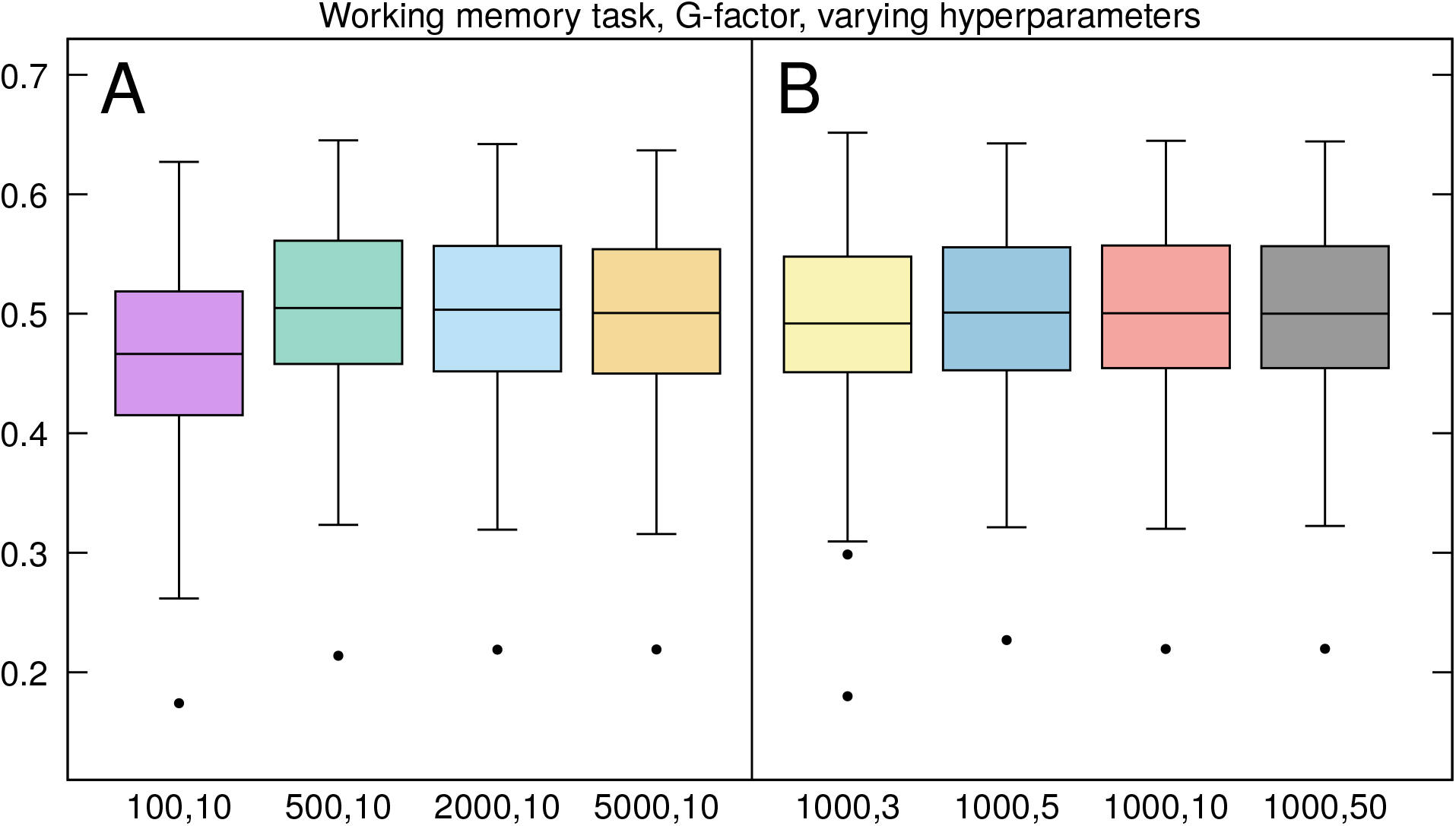
Prediction accuracy with various hyperparameter settings. Same as Supplementary Figure 12, but for the working memory task.

**Supplementary Figure 14:**
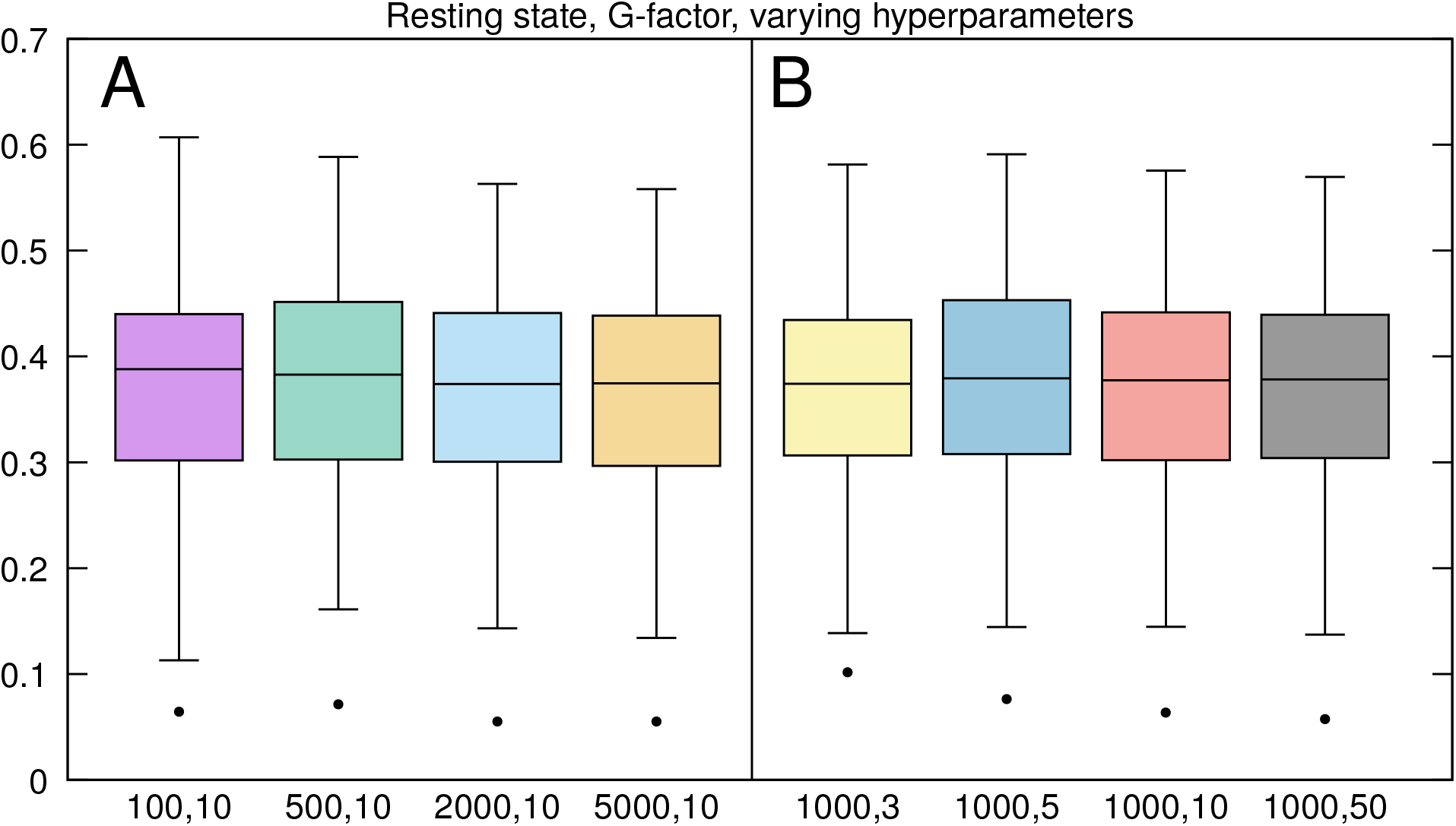
Prediction accuracy with various hyperparameter settings. Same as Supplementary Figure 12, but for resting state fMRI.

**Supplementary Figure 15:**
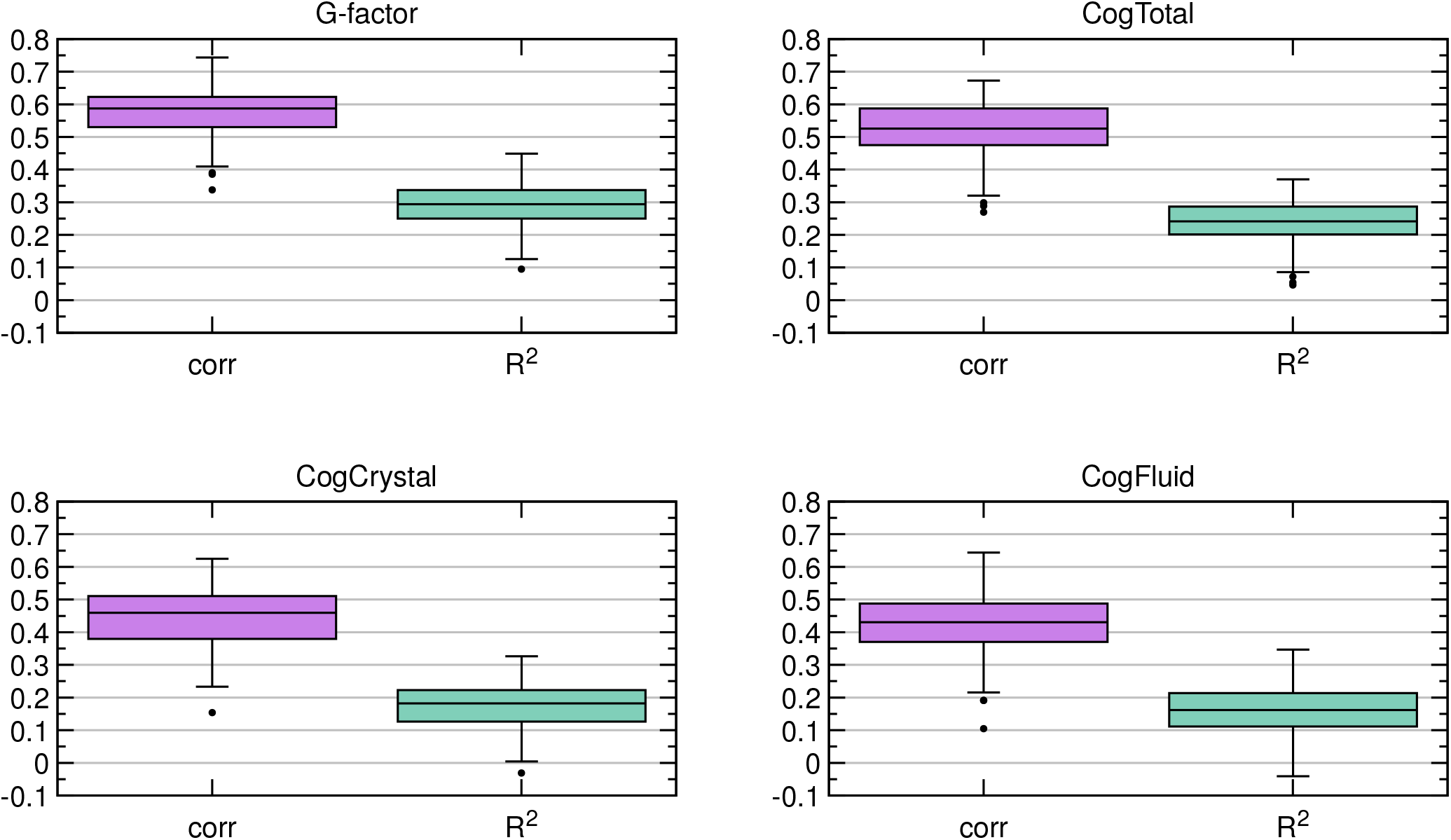
Combining two tasks. Prediction accuracy improves when the results from the two tasks (language and working memory) are combined. The boxplots show Pearson linear correlation and predictive R^2^ between observed and predicted intelligence scores.

**Supplementary Figure 16:**
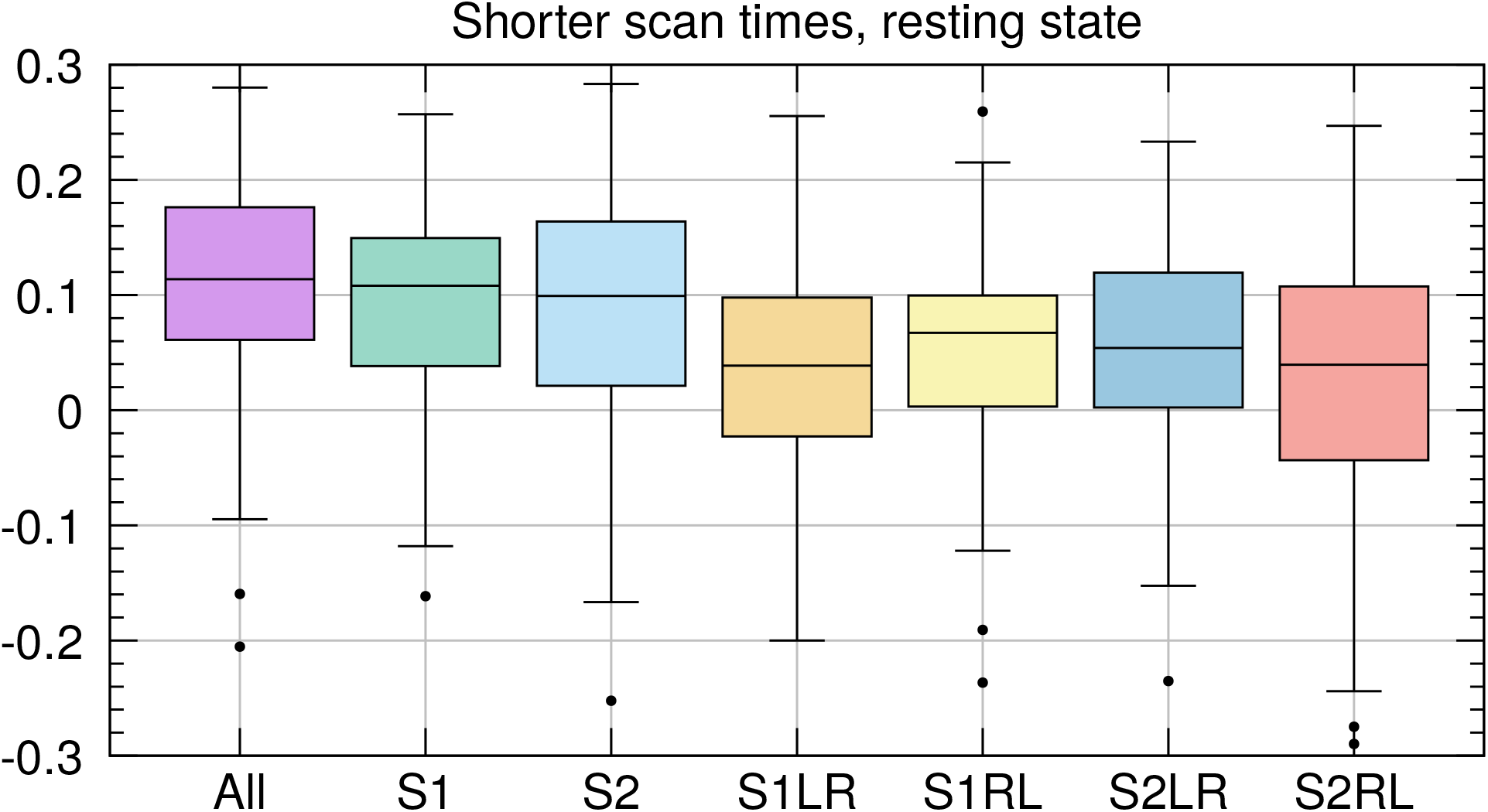
The effect of scan time on prediction accuracy (resting state). The boxplots show the reduction in predictive R^2^ as scan time is reduced. The leftmost boxplot shows the results with the complete scan time (≈ 58 min). The next two boxplots (S1,S2) show the results for session 1 and 2 (29 min each). The four rightmost boxplots show the results for each of the four runs, i.e. session 1 with LR-phase encoding, session 1 with RL-phase encoding, session 2 with LR-phase encoding, session 2 with RL-phase encoding (14.4 min each).

**Supplementary Figure 17:**
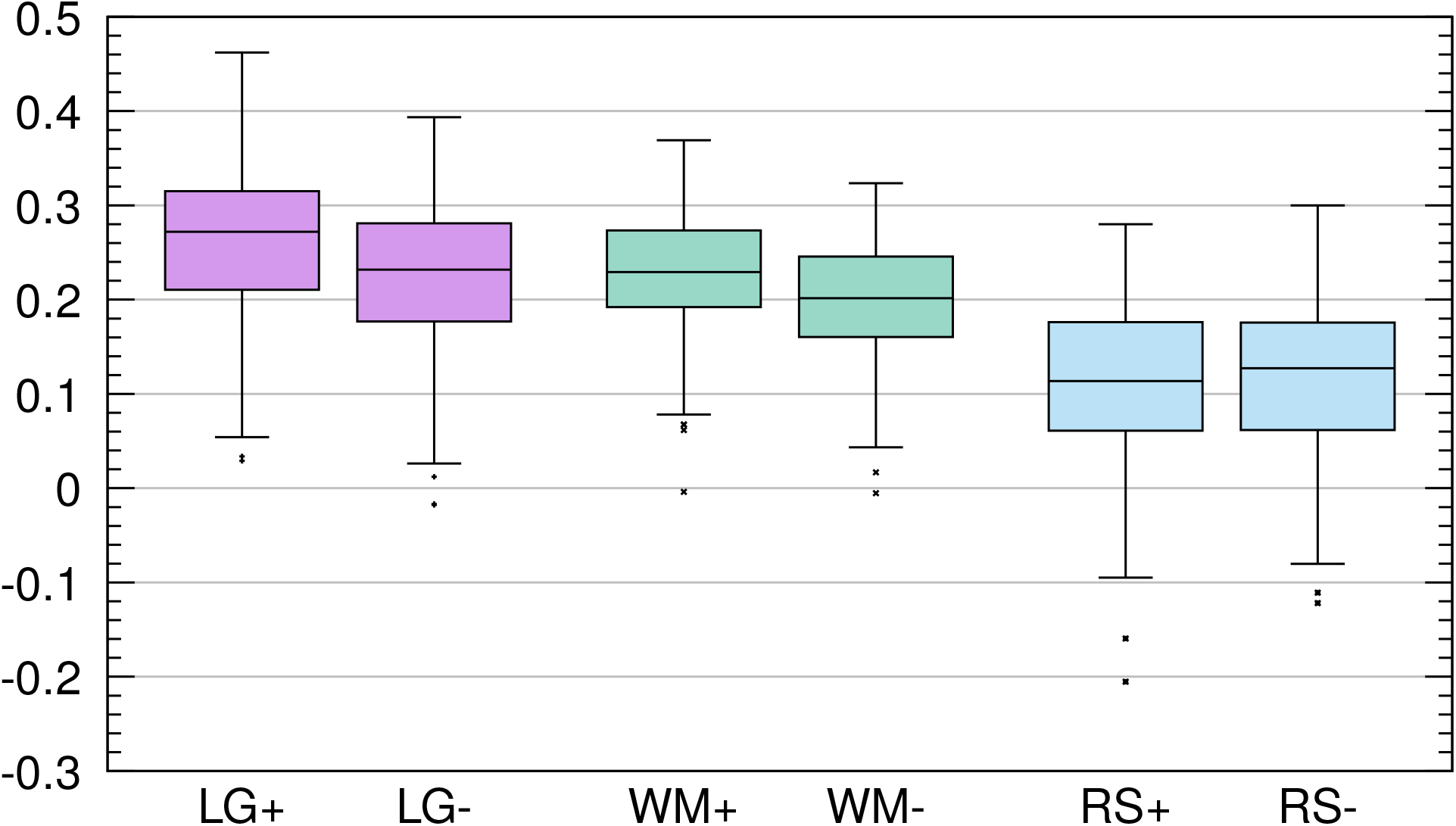
The effect of Ricci-Forman curvature maps on prediction accuracy. The thresholded curvature maps were helpful in providing more accurate predictions of the G-factor in task-fMRI, but not in resting state fMRI. The boxplots LG+ and LG-show R^2^ values for the language task with and without curvature maps, respectively. Likewise, WM+, WM-, RS+, RS-show R^2^ values for the working memory task and rs-fMRI with and without the thresholded curvature map.

**Supplementary Figure 18:**
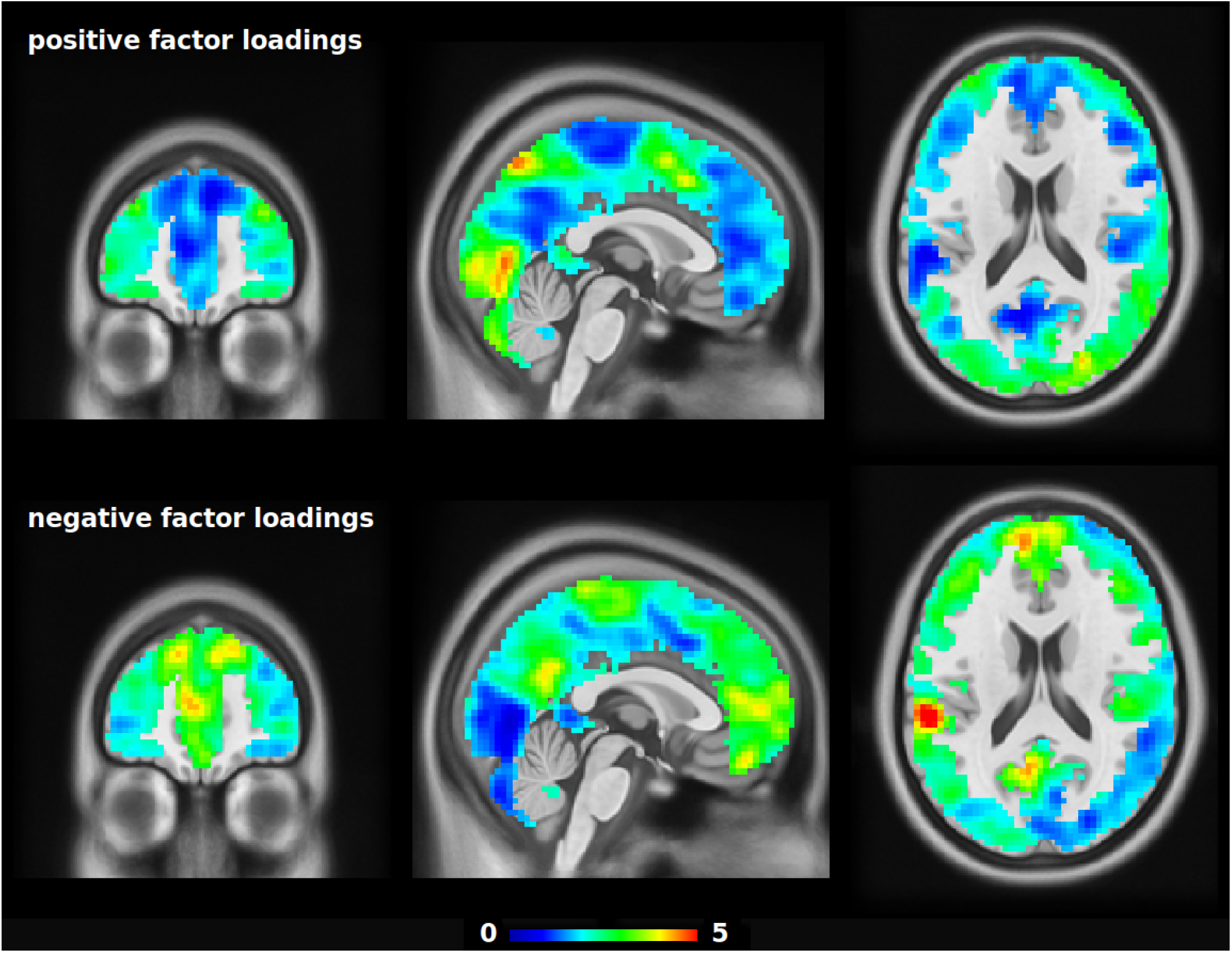
Predictive areas for general intelligence in the working memory task. The colors encode factor loadings (matrix P) estimated by partial least squares regression. Strong positive loadings indicate areas where connectivity with other brain regions is positively correlated with general intelligence. Strong negative loadings indicate areas where connectivity with other brain regions is negatively correlated with general intelligence. The colors only show relative weights, they do not have interpretable units.

**Supplementary Figure 19:**
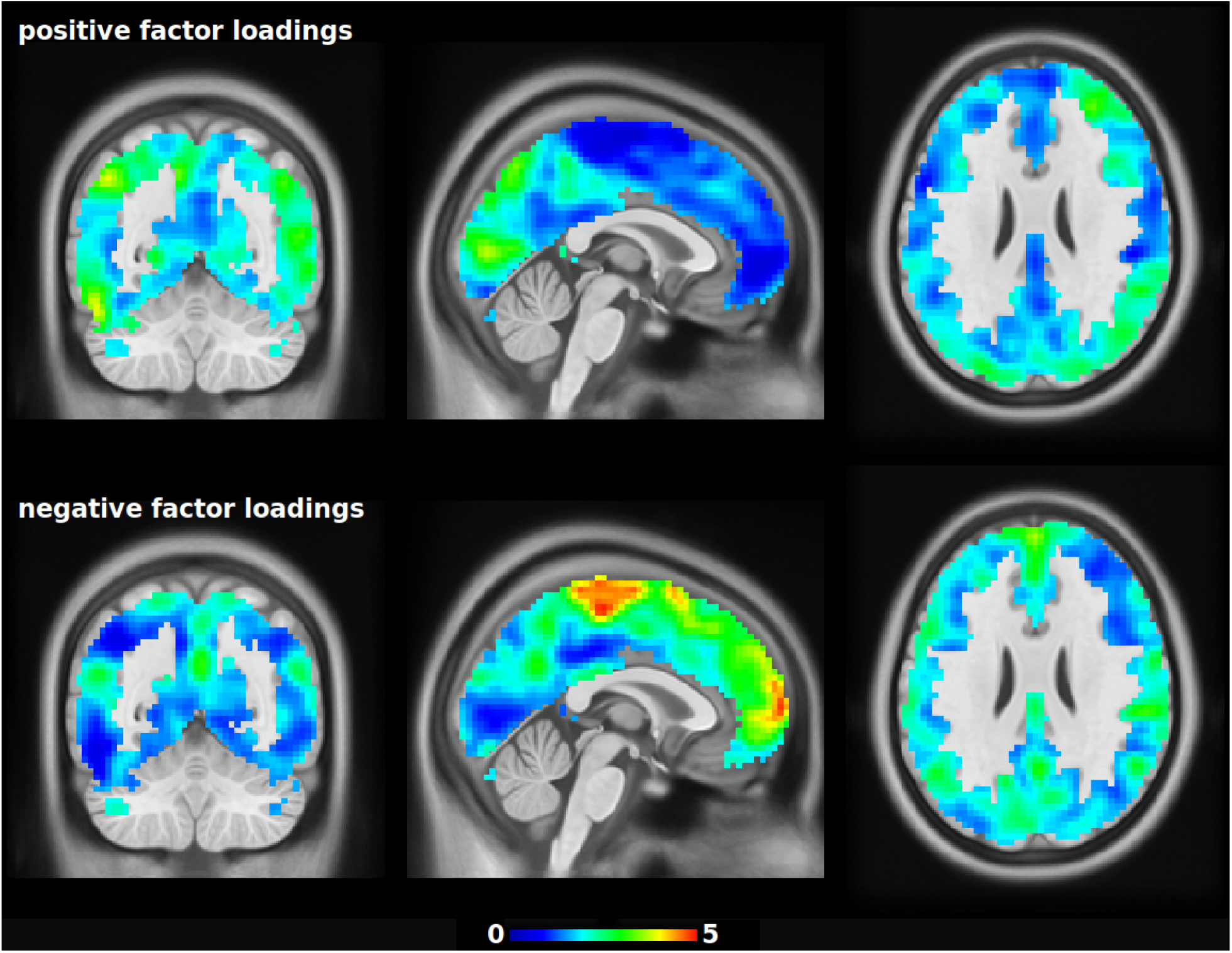
Predictive areas for general intelligence in resting state. The colors encode factor loadings (matrix P) estimated by partial least squares regression. Strong positive loadings indicate areas where connectivity with other brain regions is positively correlated with general intelligence. Strong negative loadings indicate areas where connectivity with other brain regions is negatively correlated with general intelligence. The colors only show relative weights, they do not have interpretable units.

### Row-column centering

Let *X*^*train*^ be the *n* × *m* training matrix. Let *µ* denote its total mean value, i.e.

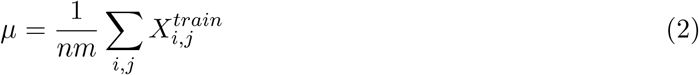

and let 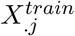 denote the mean values across column vectors *j* = 1, …, *m*, and 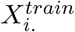 the mean values across row vectors *i* = 1, …, *n*.

Then the matrix *X*^*train*^ is centered as follows:

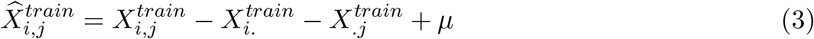

And the test matrix *X*^*test*^ is centered as follows:

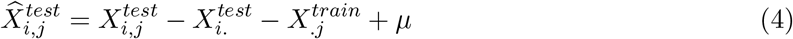

Note that the total mean *µ* and the column means 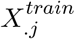, are derived from the training matrix. But the row means 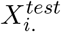 are derived from the test matrix because they correspond to individual subjects.

